# Molecular architecture and domain arrangement of the placental malaria protein VAR2CSA suggests a model for receptor binding

**DOI:** 10.1101/2020.04.16.045096

**Authors:** Maria C. Bewley, Lovely Gautam, D. Channe Gowda, John M. Flanagan

**Author notes:** Both authors contributed equally to this work (Maria C. Bewley, Lovely Gautam).

## Abstract

VAR2CSA is the placental-malaria specific member of the antigenically variant *Plasmodium falciparum* erythrocyte membrane protein 1 (PfEMP1) family. It is expressed on the surface of *Plasmodium falciparum* infected host red blood cells and binds to specific chondroitin-4-sulfate (CSA) chains of the placental proteoglycan receptor. The functional ~310 kDa ectodomain of VAR2CSA is a multi-domain protein that requires a minimum 12-mer CSA molecule for specific, high affinity receptor binding. However, how these domains interact to create the receptor binding surface is not known, limiting efforts to exploit its potential as an effective vaccine or drug target. Using small angle X-ray scattering and single particle reconstruction from negative stained electron micrographs of the ectodomain and multidomain constructs, we have determined the structural architecture of VAR2CSA. The relative location of the domains creates two distinct pores that can each accommodate the 12-mer of CSA, suggesting a model for receptor binding. This model has important implications for understanding cytoadherence of IRBCs and potentially provides a starting point for developing novel strategies to prevent and/or treat placental malaria.

## INTRODUCTION

Several species of *Plasmodium* parasites cause malaria in humans; however, the deadliest forms, including cerebral and placental malaria, are caused by *Plasmodium falciparum* (Pf) (1–3). The severity of Pf malaria is a consequence of the sequestration of parasite-infected red blood cells (IRBCs) in the vital organs of the host, leading to inflammation and fatal pathological conditions. Parasite sequestration is mediated by a family of ~60 antigenically-variant proteins, called Pf erythrocyte membrane protein 1 (PfEMP1), expressed on the IRBC surface, that are capable of binding to several host cell adhesion molecules in the vasculature of susceptible organs (4–15). Multiplication of host organ-sequestered parasites leads to microcirculatory obstruction, hypoxia, inflammation, organ dysfunction and the severe pathologies of *P. falciparum* malaria (16–21).

As a result of frequent Pf infections, most adults in malaria endemic areas have acquired protective anti-PfEMP1 antibodies to parasites strains expressing various PfEMP1s, except the placental-specific one, VAR2CSA (22). During the first pregnancy, Pf exploits the development of the placenta to overcome their pre-existing immunity by expressing VAR2CSA and sequestering in the placenta by binding to specific CSA chains of the chondroitin sulfate proteoglycan (CSPG) receptor. Subsequently, rapid parasite multiplication contributes to placental dysfunction, maternal anemia, preterm delivery, low neonate weight and maternal and pediatric morbidity and mortality (23,24); collectively these conditions are referred to as placental malaria (PM) (25–28). Women infected with Pf during prior pregnancies produce anti-VAR2CSA antibodies (29) and thus resist PM development in subsequent pregnancies (30), suggesting that VAR2CSA is a suitable therapeutic target. However, the parasite expresses polymorphic VAR2CSA, which poses challenges in the development of long-lasting efficacious vaccines against diverse CSA-binding isolates. Even so, placental sequestration of IRBCs in the intervillous spaces of the placenta and syncytiotrophoblast surface is an obligate step in the pathology of PM (31–35) that is facilitated by VAR2CSA binding to the host CSA chains of the CSPGs. Therefore, understanding the nature of the CSA:VAR2CSA interaction is a critical step toward developing effective therapeutic strategies that prevent IRBC cytoadherence in the placenta (36).

VAR2CSA is a ~350 kDa membrane protein consisting of a large ~310 kDa non-glycosylated extracellular ectodomain, a single pass transmembrane helix, and a ~42 kDa cytoplasmic acidic terminal sequence that interacts with a number of host cell and parasite-derived proteins (34). The functional CSA-binding ectodomain is comprised of six Duffy-binding-like (DBL) domains (DBL1x, DBL2x, DBL3x, DBL4ε, DBL5ε and DBL6ε) and two Interdomains (ID1 and ID2/CIDR_PAM_), connected by short linker sequences (Fig. 1A). Together, these domains form a high affinity ligand binding site with specificity for a minimum 12-mer CSA that has a characteristic, low C4 sulfate content (lsCSA) (31,33,37). In order to gain structural information, small angle X-ray scattering (SAXS) studies were performed using HEK-expressed, engineered non-glycosylated and baculovirus expressed glycosylated ectodomains (38,39). However, the resultant molecular envelopes were relatively featureless, preventing identification of individual domains or insight into carbohydrate binding. To date, there is no high-resolution structure of the VAR2CSA ectodomain, although crystal structures of *E.coli*-expressed DBL3x (40,41), DBL6ε (42,43) and the tandem DBL3x-DBL4ε domains (44) reveal a conserved fold comprised of three subdomains stabilized by multiple intra-domain disulfide bonds. Despite this, the detailed tertiary structure that forms the functional, high affinity CSA-binding surface remains unclear. It is generally agreed that domains DBL1x, ID1, DBL2x and half of ID2 (ID2a) are involved (38,39,45–47); but it is unclear whether they comprise the entire surface area for CSA binding. Therefore, to understand the details of carbohydrate binding it is essential to understand the structural architecture of VAR2CSA.

**Figure 1.**
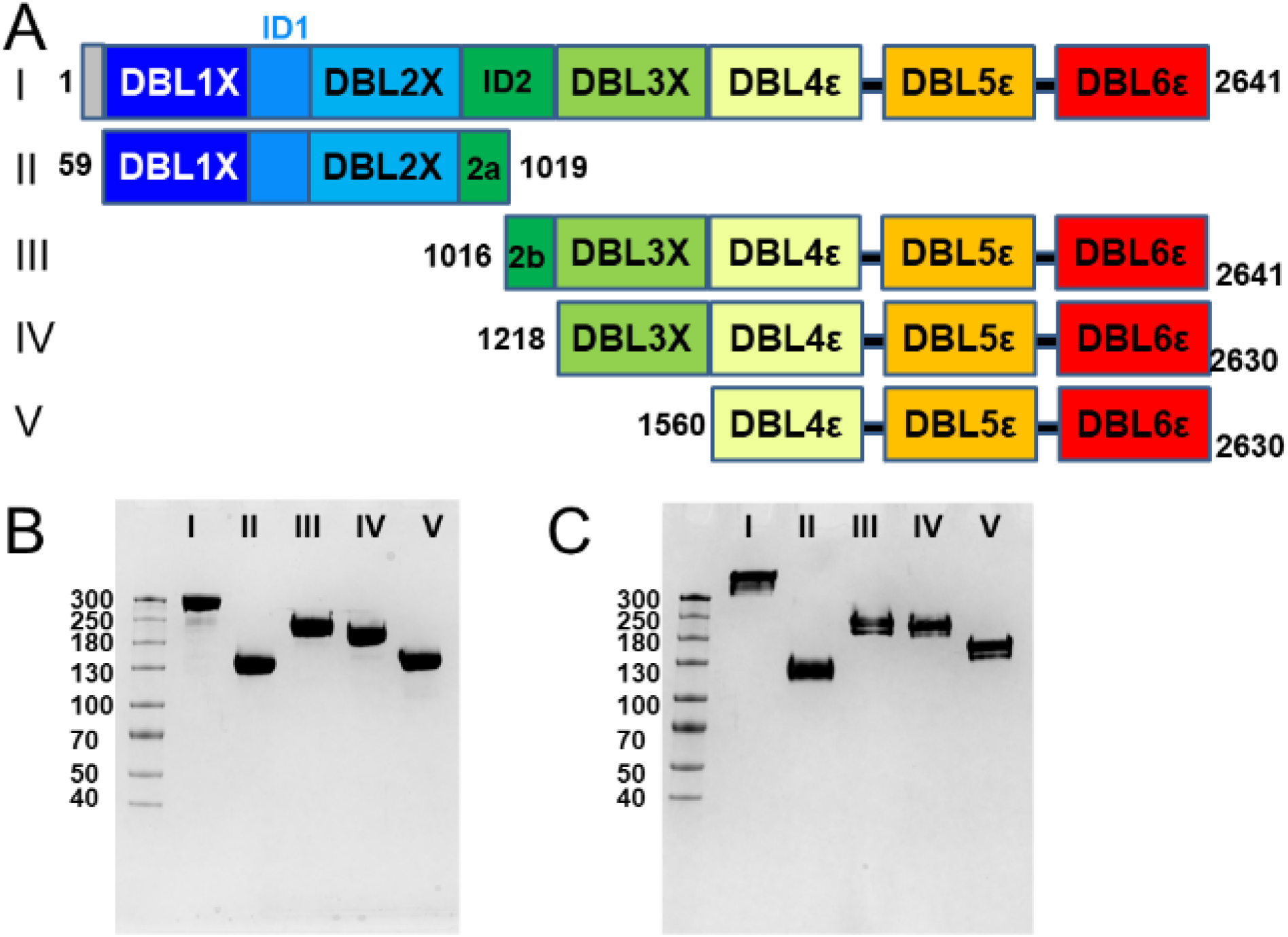
VAR2CSA and its deletion constructs expressed in HEK293-F cells and SDS-PAGE of the purified recombinant VAR2CSA proteins. (A) Schematic indicating the locations of DBL domains for constructs I-V. For each construct, the numbers at either end denote the beginning and ending amino acid sequence numbers. I, II and IV contain a non-cleavable C-terminal cMyc tag and hexahistidine tag, as coded in the commercial vector. For the III and V constructs, the C-terminal tag is replaced with a TEV-cleavable 3X-FLAG tag and a hexahistidine tag, as defined in Table S2, GBLOCK2. In addition, P1 contains a glycine and threonine residue inserted between native residues 58 and 59 as a cloning artefact. (B) Purified proteins (each 2 μg/lane), electrophoresed using 4-20 % gradient gels (1 mm thick) under reducing conditions and (C) non-reducing conditions are shown. In each gel, the lanes are: M, Molecular weight markers; I, NTS-DBL6ε (~310 kDa); II, DBL1x-ID2a (~125 kDa); III, ID2b-DBL6ε (~200 kDa); IV, DBL3x-DBL6ε (~180 kDa); V, DBL4ε-DBL6ε (~130 kDa). The molecular mass marker (kDa) standards are indicated in the left margin.

In the present study, we have expressed and characterized, structurally and functionally, the VAR2CSA ectodomain and a set of N- and C-terminal deletion constructs (Fig. 1). These proteins are folded and thermally stable and NTS-DBL6ε and DBL1x-ID2a can specifically bind the lsCSA chains of CSPG with nM affinity. Further, using a combination of size exclusion chromatography in-line with SAXS (SEC-SAXS), single particle reconstruction of negative stained electron micrographs and basic homology modeling, we have determined the relative locations of the DBL domains and produced a validated model of the VAR2CSA ectodomain. Importantly, these studies reveal, for the first time, a defined molecular shape with distinctive pores that transverse the molecule and suggest a credible model for CSA binding.

## RESULTS

### Production, purification and characterization of VAR2CSA constructs

The codon-optimized synthetic gene encoding the VAR2CSA ectodomain (NTS-DBL6ε) of *P. falciparum* 3D7 strain and four deletion constructs corresponding to DBL1x-ID2a, ID2b-DBL6ε, DBL3x-DBL6ε and DBL4ε-DBL6ε (as defined in Fig. 1A and Table S1) were expressed in HEK 293-F cells. The proteins are produced as secreted monomers and purified using a combination of ultrafiltration, nickel-affinity, cation-exchange, and size-exclusion chromatography. After purification, a single band, at the expected apparent molecular mass, is observed upon SDS-PAGE under reducing conditions (Fig. 1B). Similarly, under non-reducing conditions a single band is observed (Fig. 1C). The purity of each protein construct, as assessed by the SDS-PAGE band profiles, is ~98%. Yields of the purified recombinant proteins range from 0.5-0.7 mg/l for NTS-DBL6ε to 14 mg/l for DBL4ε-DBL6ε. Several additional deletion constructs (DBL1x-DBL5ε, DBL1-ID2b, ID1-DBL6ε, ID1-ID2) were tested for protein production in our system, but these either failed to express or to purify as folded monomers and thus were not analyzed further.

To assess the folding of the purified recombinant proteins, far UV CD spectra were recorded at 25 °C and evaluated. A strong maximum at 200 nm and troughs at 208 nm and 222 nm are observed (Fig. S1A) that are characteristic of proteins containing substantial α-helical secondary structures. The spectra are similar to results reported for VAR2CSA constructs DBL6ε, DBL4ε-DBL6ε and DBL1x-DBL6ε (38). The stability of each construct was investigated by following the changes in ellipticity at 222 nm as a function of temperature between 25-85 °C. In all cases, a transition occurs between 70 and 75 °C (Fig. S1B), demonstrating that these constructs are thermally stable. The observed transition temperatures are similar to those reported earlier for analogous constructs (38). However, for each construct, the spectrum collected at 90 °C still shows residual ellipticity at 208 nm and 222 nm (Fig. S1C-G), demonstrating that unfolding is incomplete, preventing a thermodynamically-justified value for T_m_ from being calculated. Indeed, the complete unfolding of NTS-DBL6ε is achieved only in the presence of the reducing agent tris(2-carboxyethyl) phosphine (Fig. S1H). These results are consistent with the constructs being stable, as expected for domains containing a large number of disulfide bonds (48).

The binding of each VAR2CSA construct to CSPG fraction II (CSPG-II) purified from bovine cornea (49), which contains chondroitin sulfate chains having low levels of 4-sulfate groups, was assessed in an ELISA-based assay (Fig. S2A). NTS-DBL6ε binds with a high affinity (K_D_ 19 ± 1 nM), whereas DBL1x-ID2a binds with a much lower affinity (K_D_ 113 ± 24 nM). No other deletion construct (ID2b-DBL6ε, DBL3x-DBL6ε and DBL4ε-DBL6ε) show measurable binding in this assay. Competition studies with NTS-DBL6ε and DBL1x-ID2a using bovine tracheal CSA and CSC, which contain different relative proportions of CSA, show that CSA efficiently inhibits the binding of both constructs to CSPG-II. CSC, which contains 10-20 % CSA and 80-90 % C6S, shows relatively lower inhibition. HA, a polymer of disaccharides composed of D-glucuronic acid and *N*-acetyl-D-glucosamine and no 4-sulfated groups, also has no measurable inhibition (Fig. S2B). Together, these results demonstrate that the purified NTS-DBL6ε and DBL1x-ID2a bind low sulfated CSA and that the former binds with higher affinity than the latter.

### SAXS analysis suggests that the VAR2CSA ectodomain is compact

In order to gain insight into the structural organization of the ectodomain, NTS-DBL6ε and the deletion constructs, DBL1x-ID2a, DBL3x-DBL6ε and DBL4ε-DBL6ε, were characterized using SEC-SAXS (Table 1, Figs. 2, S3). The data were processed using the ATSAS Suite (50,51) and the values obtained from this analysis compared to those previously reported, if available (Table 1 and Figs. 2A-D, S3). All of the constructs are linear in the Guinier region of the SAXS curve over the given qR_g_ ranges (Fig. 2E), indicating that they are monodisperse. The R_g_ value obtained for DBL1x-ID2a is smaller than that reported previously (39), which is consistent with the removal of small amounts of dimer and higher order aggregates during chromatography and the absence of glycosylation in our constructs. It should be noted that, although the exact domain boundaries are slightly different (Fig 1A), this modification alone would not account for the observed differences in R_g_ values. For each construct, assessment of the molecular weight (Table 1) using a consensus Bayesian method, implemented in ATSAS 3.0 (52), is in excellent agreement with both the theoretical value and those obtained by SDS-PAGE, providing further support that each of the proteins exist as a monomer in solution. For NTS-DBL6ε, DBL1x-ID2a, and DBL3x-6ε, the pair-distribution, P(r), function approaches zero smoothly at r = 0 Å and the D_max_ and has a single peak consistent with relatively globular molecules (Fig. S3A). By contrast, the P(r) function for DBL4ε-DBL6ε contains an asymmetric peak with a maxima at ~42 Å and a shoulder centered at ~62 Å, indicative of a molecule that contains spatially distinct modules, with the first peak due to the distribution of scattering centers (chords) within a module and the shoulder reflecting the chords between the modules. The dimensionless Kratky plots further support this analysis (Fig. S3B). NTS-DBL6ε and DBL1x-ID2a have similar plots that contain a single peak at qR_g_ = ~1.7 and (qR_g_)^2^I(q)/I(0) = 1.1, in agreement with theoretical values of 1.75 and 1.1 for a globular protein (53), respectively, and returning to zero at qR_g_ ~5 (Fig. S3B). The location of the peak is consistent with each construct behaving in solution as a single module. By contrast, the plots for the DBL3x-DBL6ε and DBL4ε-DBL6ε contain peaks that are shifted to longer qR_g_ (1.8-1.95), indicative of a more elongated molecule. In addition, the plots for DBL3x-DBL6ε and DBL4ε-DBL6ε contain a shoulder at qR_g_ ~4.9 and qR_g_ ~4 respectively, and return to zero at qR_g_ ~10, characteristic of proteins containing spatially distinct modules (54).

**Table 1.** Data Processing Statistics from SAXS measurements.

| | R<sub>g</sub> | qR<sub>g</sub><br>range for<br>Guinier | Io§ | D<sub>MAX</sub><br>from<br>P(r) | Molecular<br>weight<br>estimate*# | Credibility<br>interval# | NSD<br>(SD) | $\chi^2$<br>CorMap<br>p-value | Ambiguity |
| --- | --- | --- | --- | --- | --- | --- | --- | --- | --- |
| NTS-DBL6ε | 55 | 0.5-1.3 | 104.1<br>(0.7) | 177 | 318450<br>(86 %) | [221050, 372700]<br>(100 %) | .81<br>(0.03) | 1.1<br>(0.1) | 1.18 (5) |
| DBL1x-DBL6ε<sup>a</sup> | 62 | NP |  | 185 | NP | NA | NP | NA | NA |
| DBL1x-DBL6ε<sup>b</sup> | 63 | 0.8-1.3 |  | 219 | 360/431 | NA | .72-.8<br>(NP) | 1.1<br>(NA) | NA |
| DBL3x-DBL6ε | 53 | 0.4-1.3 | 338.5<br>(1.0) | 160 | 208000<br>(84 %) | [176600, 221050]<br>(100 %) | 0.95<br>(0.37) | 0.9<br>(0.4) | 2.31 (5) |
| DBL4ε-DBL6ε | 49 | 0.6-1.3 | 22.1<br>(0.1) | 148 | 157050<br>(68 %) | [151450, 176600]<br>(99 %) | 0.98<br>(0.04) | 0.8<br>(0.1) | 2.42 (5) |
| DBL1x-ID2a | 38 | 0.4-1.3 | 12.1<br>(0.3) | 132 | 94225<br>(37 %) | [77400, 99200]<br>(91 %) | 0.99<br>(0.28) | 0.5<br>(0.5) | 1.57 (5) |
| DBL1x-ID2a<sup>b</sup> | 48 | 0.9-1.3 |  | 166 | 167/153 | NA | .72-.8<br>(NP) | 2.5<br>(NA) | NA |
| DBL1x-DBL4ε<sup>b</sup> | 49 | 0.6-1.3 |  | 169 | 207/249 | NA | .72-.8<br>(NP) | 1.5<br>(NA) | NA |
| ID1-DBL4ε<sup>b</sup> | 55 | 0.8-1.3 |  | 188 | 307/483 | NA | .72-.8<br>(NP) | 1.3<br>(NA) | NA |

NP - not published; NA - not available at the time of their publication.
§Number in parentheses are SD.
\*Numbers in parentheses are probability estimates of the molecular weight.

**Figure 2.**
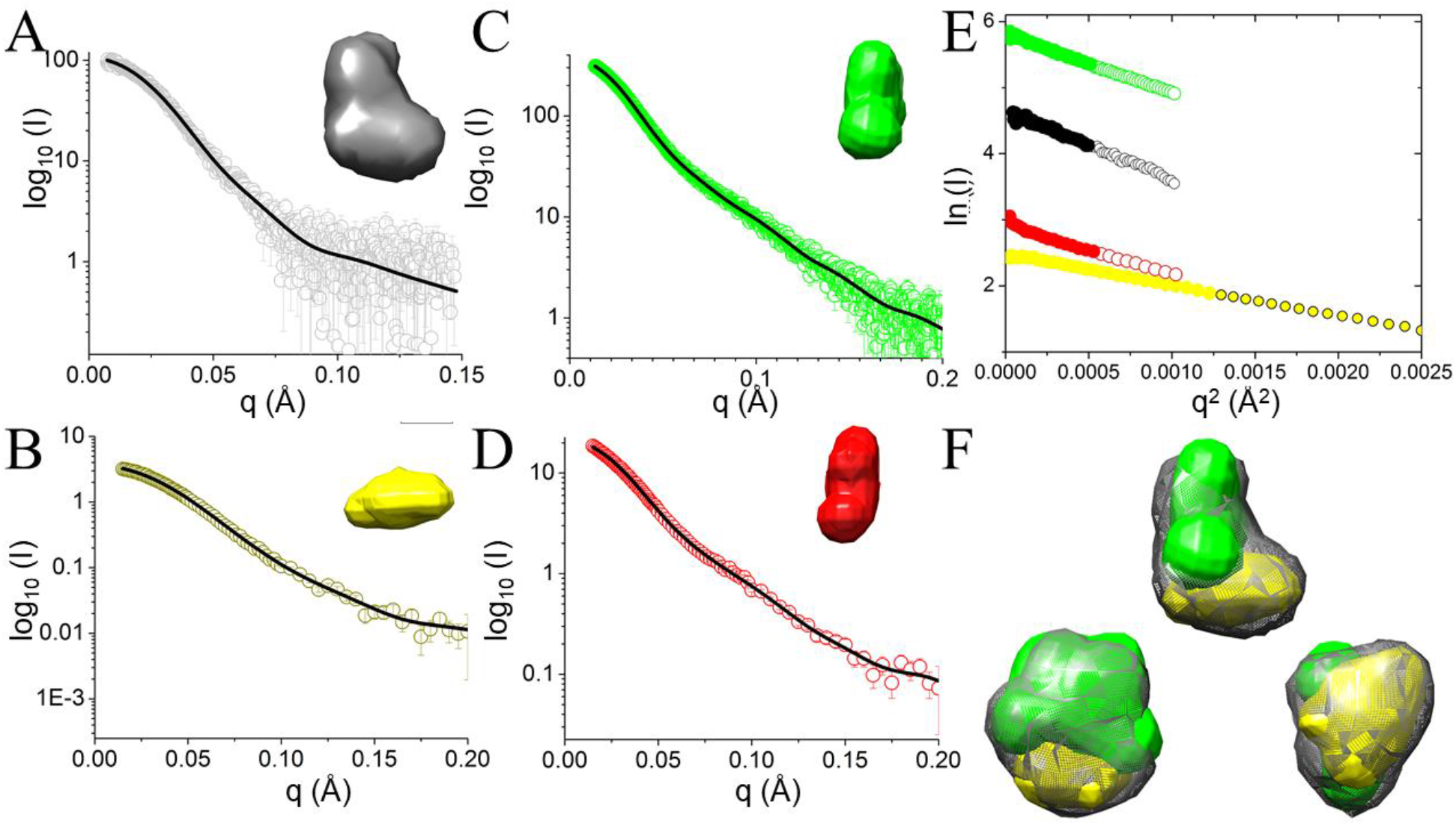
SAXS analysis indicates that purified VAR2CSA constructs are asymmetric and compactly folded. (A-D) Experimental scattering intensities (open circles) and calculated fit (black lines) as a function of the momentum transfer (q) for constructs with a single view of the averaged ab initio envelope(inset) (A) NTS-DBL6ε, black; (B) DBL1x-ID2a yellow; (C) DBL3x-DBL6ε, green; (D) DBL4ε-DBL6ε red. (E) Guinier plot showing the fitted region as large filled circles (linear fit model; Pearson CC −0.99 for all constructs) and smaller open circles indicating data beyond the Guinier region, colored as in A-D. (F) Three orthogonal views of the *ab initio* envelopes of NTS-DBL6ε (grey mesh) reveal an asymmetric molecule with a larger broad base and smaller, narrower top. *ab initio* envelopes for DBL1x-ID2a (yellow) and DBL3x-DBL6ε (green) show that the envelopes of the two deletion constructs account for the NTS-DBL6ε envelope within the limits of the resolution. For each construct, the averaged/filtered *ab initio* bead models were converted to a pseudo surfaces using Chimera (55).

### Ab initio bead model of VAR2CSA constructs reveal their overall shape

For each construct, an *ab initio* averaged/filtered bead model was calculated from the models of 20 individual simulations with DAMMIF in ATSAS 3.0. The simulated SAXS curves for these models are in good agreement with the data, as judged by the χ2 values (Table 1). For NTS-DBL6ε, the ensemble of individual calculated models are similar (Table 1), resulting in an asymmetric bead model of approximate dimensions 160 Å × 130 Å × 90 Å. It has a thinner, ~ 50 Å wide, head in its narrowest dimension and a wider base of ~ 90 Å (Fig. 2A). This is in good agreement with the approximate dimensions of crystal structures of DBL domains; for example, the crystal structure of DBL6ε (42) is an elongated molecule of approximate dimensions 35 Å × 45 Å × 85 Å (calculated R_g_ 23 Å) that can be accommodated within either the head or the base.

In order to identify the location of DBL1x-ID2a, DBL3x-DBL6ε and DBL4ε-DBL6ε within NTS-DBL6ε, *ab initio* bead models were calculated as before. For DBL1x-ID2a, the averaged bead model is smaller than NTS-DBL6ε, as expected from its smaller molecular weight, with approximate dimensions of 115 Å × 75 Å × 50 Å (Fig. 2B). For DBL3x-DBL6ε, the averaged model is asymmetric and elongated in shape (135 Å × 100 Å × 75 Å) (Fig. 2C). For DBL4ε-DBL6ε, the consensus bead model is a triangular prism reminiscent of a letter T (165 Å × 100 Å × 75 Å) with the vertical leg ~ 50 Å in diameter and the horizontal arm approximately 115 Å × 65 Å × 45 Å (Fig. 2D).

The bead models of the fragments were then fit into NTS-DBL6ε using chimera, placing DBL3x-DBL6ε first because it was the largest fragment. Although individually, each could occupy a range of positions, there was only one arrangement in which DBL3x-DBL6ε and DBL1x-ID2a fit into the full-length envelope simultaneously without a large conformational rearrangement (Fig. 2F). DBL4ε-DBL6ε fits inside DBL3x-DBL6ε leaving an additional volume that, presumably, corresponds to DBL3x; however, it could not be placed in a unique orientation. Thus the SAXS data are consistent with DBL1x-2IDa occupying the base of the molecule and DBL3x-DBL6ε occupying the length of the molecule including a thinner head region.

### EM analysis of negatively stained single particles reveal the spatial arrangements of DBL domains in VAR2CSA

In order to generate a higher resolution model than possible by SEC-SAXS analysis, single particle reconstruction of negative stained images was calculated for all of the constructs. During data collection, examination of regions throughout the grids revealed that all constructs produced both positively and negatively stained images on the same grid and, in some cases, within the same field. Therefore, data were collected from areas of the grids in which negatively stained particles dominated. Fig. 3A shows a representative negative-stained micrograph with well isolated NTS-DBL6ε particles. 2D classification of particles picked from this and other equivalent images (Fig. 3B; S4) gives a range of well-defined classes that, in side-view, suggests a roughly rectangular particle with three layers of stain-excluded density (L1-3) arranged into two asymmetrically-distributed lobes (Fig. 3C). The smaller lobe (L1) contains a single distinct volume and is referred to hereafter as the head, whereas the larger lobe (L2, L3) is comprised of three distinct regions, hereafter called the body, the tail and the feet. Further, inspection of these side-on classes suggest that the head is connected to the body via a thin, stain-excluding neck (red arrows in Fig. S4). A second, smaller stain-excluding volume from the middle of the head to the body (green arrows in Fig. S4) may also be a direct connection or, alternatively, result from a superposition of distinct structural features in some projections. The location of the C-terminus of DBL6ε in the structure was identified in negative stain images of NTS-DBL6ε bound to an anti-cMyc IgG type monoclonal antibody (mAb; Fig. 3D). Particles were picked from this and equivalent images to obtain 2D-classes which fit into one of three distinct subsets: classes similar to those observed with NTS-DBL6ε alone (Fig. 3E); Y-shaped classes in projection that are characteristic views for the free mAb, (Fig. 3F); and classes corresponding to side-on images of complexes between NTS-DBL6ε and the Y-shaped mAb (Fig. 3G). In this final set of 2D-classes, the Y-shaped mAb is clearly visible at the end of the head, providing an unambiguous identification of the location of the C-terminal cMyc tag adjacent to the head domain (small lobe). It follows that DBL6ε is located in the density adjacent to the bound antibody at the free end of the head, furthest away from the neck and body. The relatively short linker between DBL5ε and DBL6ε (~10-20 amino acids) and the clear connectivity of the density in the head suggests the remaining head density arises from DBL5ε.

**Figure 3.**
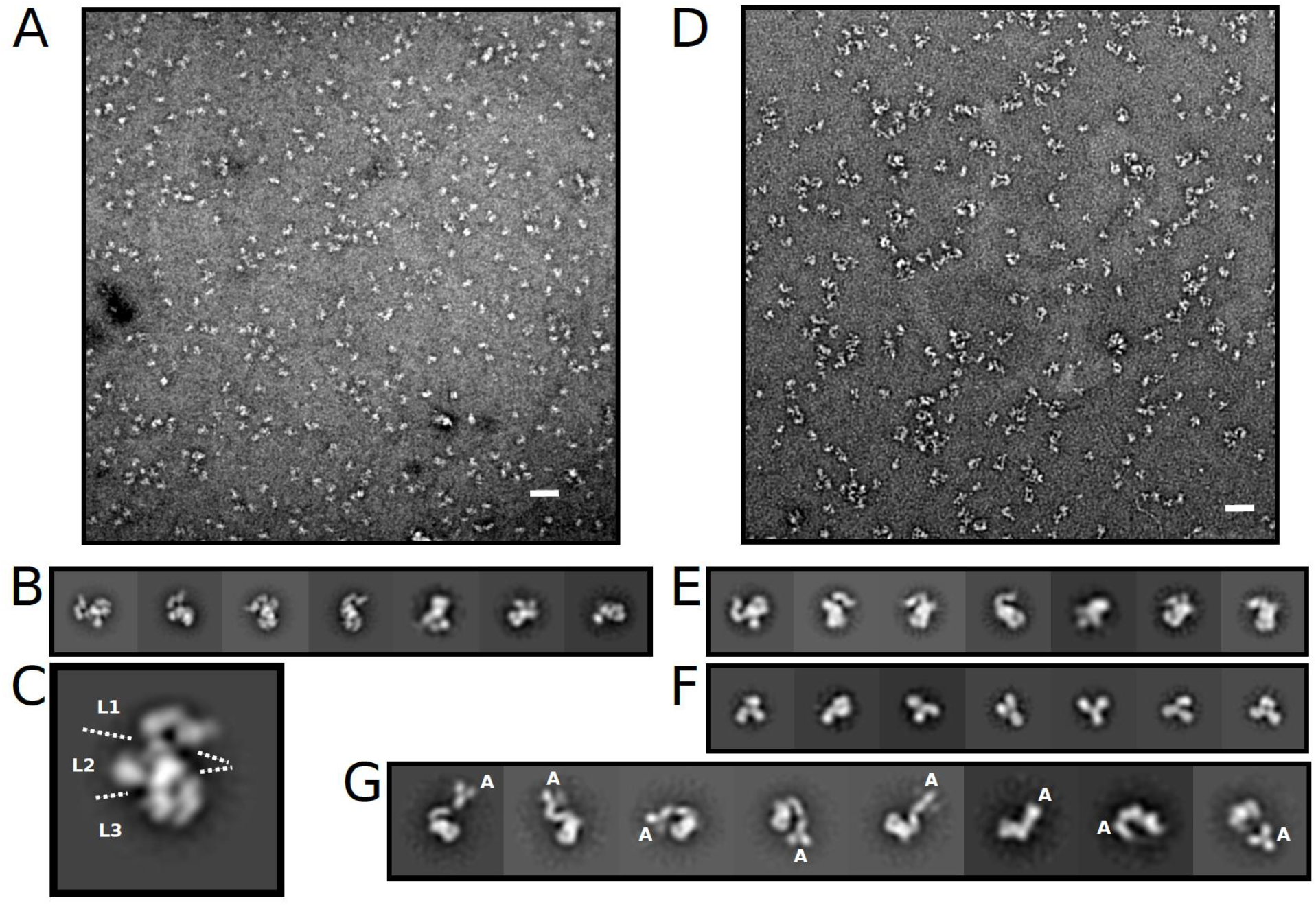
VAR2CSA is an asymmetric molecule with the C-terminus located within the smaller lobe. (A) Micrograph of NTS-DBL6ε. The white bar is 50 nm. (B) Subset of 2D classes obtained during initial 2D classification of NTS-DBL6ε. (C) Single 2D class showing the layers (L1, L2, L3), separated by broken white lines, defined in the text. (D) Micrograph of NTS-DBL6ε bound to anti-cMyc antibody. (E) Representative 2D classes of unbound NTS-DBL6ε. (F) Representative 2D classes of unbound anti-cMyc antibody. (G) Representative 2D classes of NTS-DBL6ε:anti-cMyc-antibody complex. The letter A indicates the location of the cMyc antibody at the tip of the smaller lobe of NTS-DBL6ε.

Particles selected from the best 2D classes were used to reconstruct the 3D volume of NTS-DBL6ε (Fig. 4A; Table 2). This reconstruction shows that the ectodomain adopts a slightly extended conformation with overall dimensions 175 Å × 160 Å × 110 Å (Fig. 4A) that is slightly larger than those estimated from the corresponding SAXS bead model. Consistent with the views in the 2D classes, the 3D volume is comprised of three layers that are together broadly reminiscent of a duck, with a head, body, feet and a tail (Fig. 4A). The strong concordance between the reprojections from the 3D volumes and their corresponding 2D classes (Fig. 4B; Table 2) provides validation for the NTS-DBL6ε model. The first layer comprises the head (125 Å × 75 Å × 65 Å), and is of sufficient volume to accommodate two DBL domains. The middle layer has a larger volume comprising the body (100 Å × 80 Å × 60 Å) and bulbous DBL domain-sized tail (65 Å × 60 Å × 55 Å) that protrudes from it. The final layer is a C-shaped volume (120 Å × 85 Å × 60 Å) comprising the feet; a central cleft (20 Å × 30 Å × 95 Å) separates each DBL sized foot.

**Table 2.**
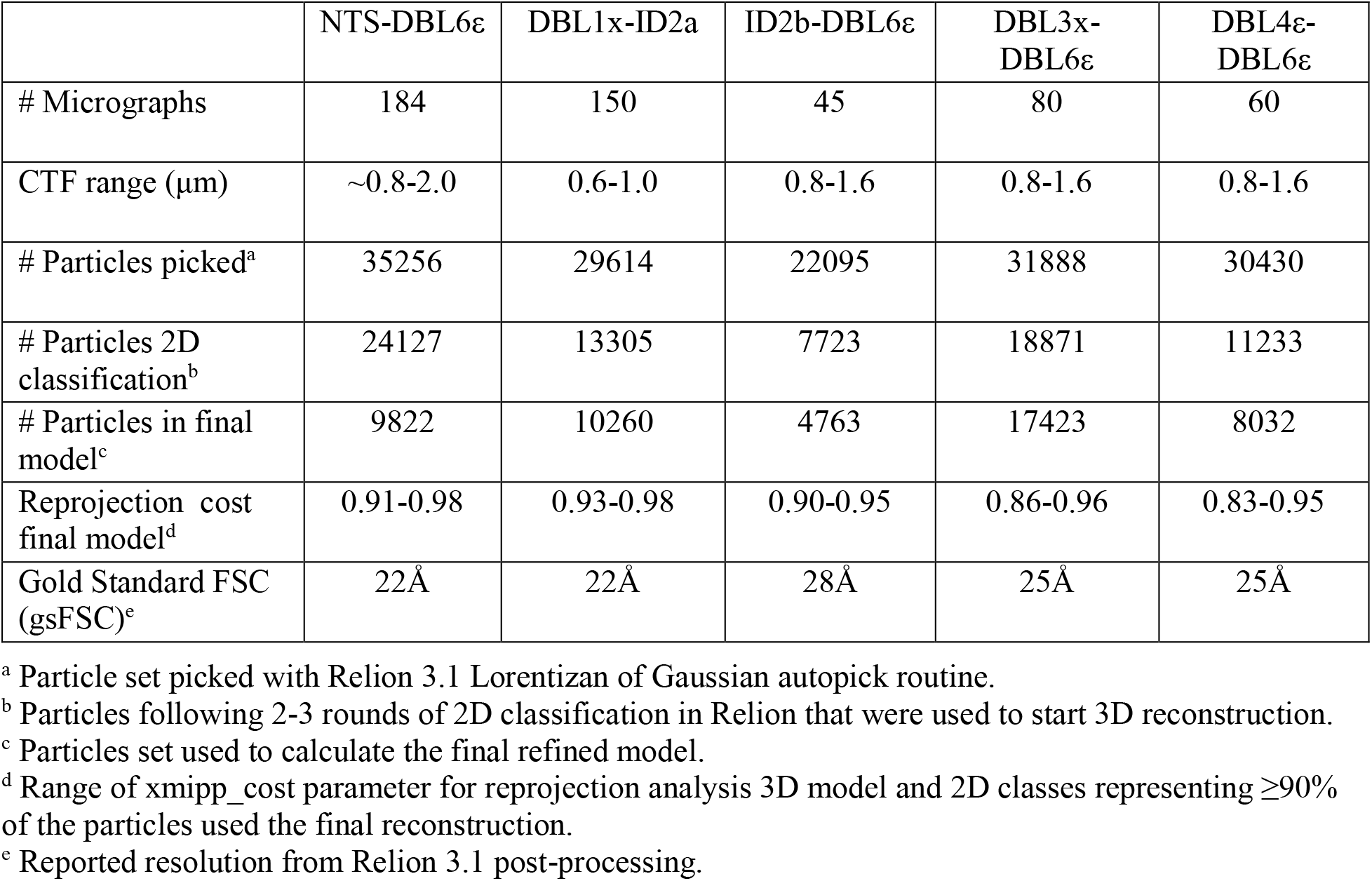
Data processing statistics for single particle reconstruction of negative stained EM.

**Figure 4.**
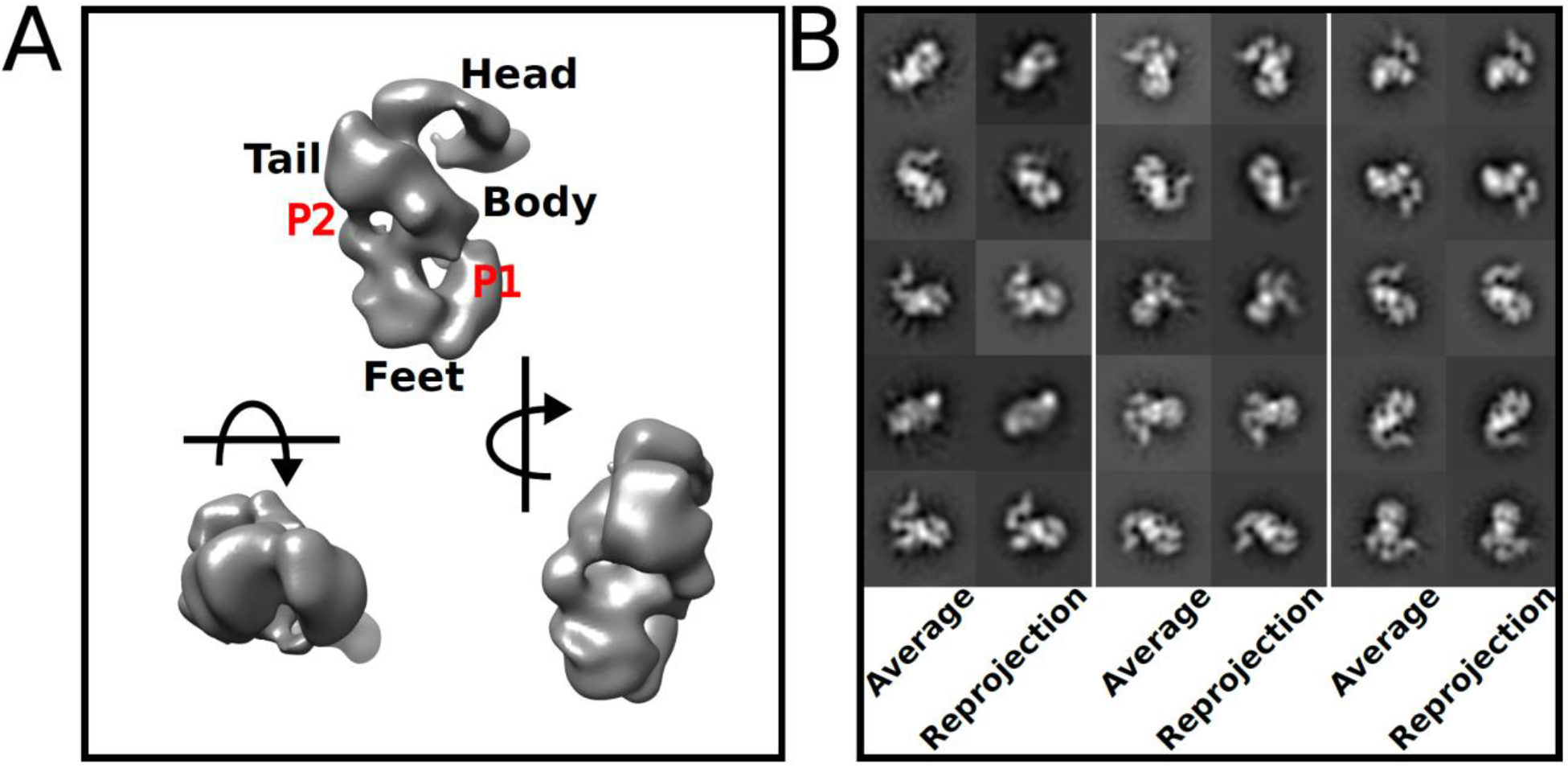
The NTS-DBL6ε reconstructed volume contains distinct regions and is consistent with the 2D classes. (A) Three orthogonal views of the 3D reconstructed NTS-DBL6ε molecule with head, tail, body and feet, as indicated. Two pores P1 and P2 run through the protein. The arrows show the direction of rotation relative to the top left view (B) The top 15 2D classes, representing ≥ 90 % of the particles used in the 3D reconstruction, paired with their corresponding back projection of the 3D model. All 3D volumes in this and subsequent figures were visualized using Chimera (61).

To determine whether the particular structural features in layers L1-L3 of NTS-DBL6ε could be assigned to specific regions of the protein, single particle reconstructions of negative stained EM images of the N- and C-terminal deletion constructs ID2b-DBL6ε and DBL1x-2IDa, respectively, were calculated and the resulting volumes compared to the EM-based reconstruction of NTS-DBL6ε. For the N-terminal deletion construct ID2b-DBL6ε (Fig. 5A, S5), an S-shaped molecule containing two weak stain-excluding regions was visible in some 2D classes (denoted by red and green arrows in Fig. S5A) strongly resembling the L1 and L2 regions observed in 2D classes of NTS-DBL6ε. This S-shaped arrangement also defined the 3D volume which is comprised of two cylindrical arms ~60 Å in diameter and 120 Å in length (Fig. 5A) and had similar characteristics to the head, tail, and body volumes of NTS-DBL6ε. For the C-terminal deletion construct, DBL1x-ID2a, an asymmetric bilobal molecule, was visible in 2D classes most strongly resembling the L3 region observed in the 2D classes of NTS-DBL6ε. The corresponding reconstructed volume is globular (125 × 85 × 80 Å) with a distinct cleft approximately 20 Å in diameter (Fig. 5B, S6) that is also visible in the feet of the NTS-DBL6ε volume. Attempts to identify the location the C-terminal cMyc tag in the DBL1x-2IDa volume were unsuccessful (Fig. S7) preventing specific assignment of DBL1x and DBL2x domains. The assignment of 3D features was confirmed by superposition of the DBL1x-ID2a and ID2b-DBL6ε volumes into that of NTS-DBL6ε (Fig. 5C, D): As expected from our analysis, the DBL1x-ID2a volume localizes to the ‘feet’ and ID2b-DBL6ε to the head, body and tail features of the NTS-DBL6ε volume, respectively. Together, these data demonstrate that L1 contains DBL5ε and DBL6ε; L2 contains DBL3x and DBl4ε; L3 contains DBL1x, ID1 and DBL2x. Placed in this manner, the tip of the larger density in the DBL1x-ID2a reconstruction extends from the feet into the region in the body near the DBL3x-DBL4ε volume. This suggests that DBL2x and ID2a would be contained in the larger volume and thus DBL1x in the smaller volume of the DBL1x-2IDa reconstruction.

**Figure 5.**
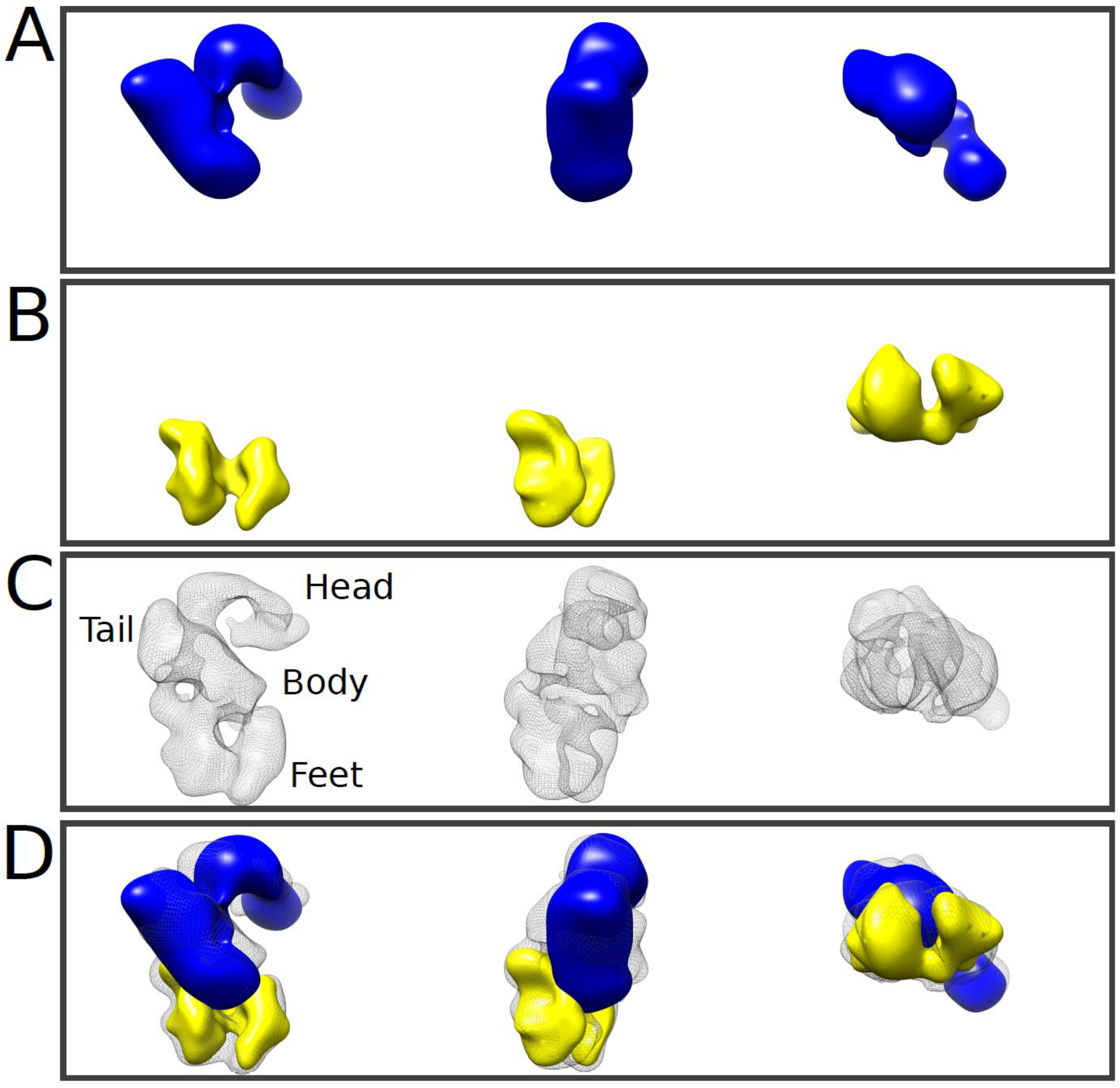
The reconstructions delineate the locations of the DBL domains. Three orthogonal views of the reconstructed volume of (A) ID2b-DBL6ε (blue solid) (B) DBL1x-ID2a (yellow solid) (C) NTS-DBL6ε (black mesh). (D) A-C superimposed. ID2b-DBL6ε, forms the head and tail of the molecule and DBL1x-ID2a the feet. ID2 appears to form a bridge linking DBL1x and DBL2x to the remainder of the molecule.

To further assign volumes in L2, single particle reconstructions of DBL3x-DBL6ε (Fig. S8) and DBL4ε-DBL6ε (Fig. S9) were calculated and aligned with that of ID2b-DBL6ε (Fig. 6A). The DBL3x-DBL6ε construct (Fig. 6A; S8) is an S-shaped molecule (approximate dimensions 160 Å × 135 Å × 65 Å) with four clear, stain excluding regions in both 2D classes and in the 3D reconstructed volume. The DBL4ε-DBL6ε construct is a skewed T-shaped molecule (approximate dimensions 150 Å × 130 Å × 65Å), with 3 clear stain excluding volumes which is evident in both the 2D classes and in the 3D map (Table 2 and Fig. 6A, Fig. S9). The relatively weaker stain excluding densities connecting L1 and L2 observed in ID2b-DBL6ε, are less striking in the DBL3x-DBL6ε and DBL4ε-DBL6ε reconstructions, however, corresponding maps and projections are consistent with the observed 2D classes (Fig. S8). The weaker connecting density is likely due to a combination of a lower signal-to-noise ratio of images of these constructs and, potentially, relatively more conformational freedom (Table 2). The volumes of DBL3x-DBL6ε and DBL4ε-DBL6ε were fit into that of ID2b-DBL6ε using an automated fitting algorithm in Chimera (Fig. 6A). Since the distinctive volume of DBL5ε and DBL6ε in L1 is common to all N-terminal deletion constructs, it provided a visual constrain for assessing fit. In each case, applying this constrain, the placement of volumes is unique and reveals that in L2, DBL4ε is located below both DBL5ε and DBL6ε in the body and DBL3x forms the tail (Fig. 6A). It should be noted that for all constructs containing DBL4ε, DBL5ε and DBL6ε, the domains occupy equivalent locations in all maps, however, in the DBL3x-DBL6ε map, the volume corresponding to DBL6ε is rotated by ~45 ° relative to its location in NTS-DBL6ε and ID2b-DBL6ε. The distinctly different orientation of the terminal domain is also evident when comparing some 2D classes (Fig. 6B, bottom panel). Further, although the two arms of the S-shape are better defined in the volume of ID2b-DBL6ε (Fig. 6A), there was no obvious connected density that could be assigned to ID2b. It is not clear whether the presence of ID2b directly improves the reconstruction quality via a stabilization of the structure or it is simply due to the higher signal-to-noise of the negative stained images for this construct.

**Figure 6.**
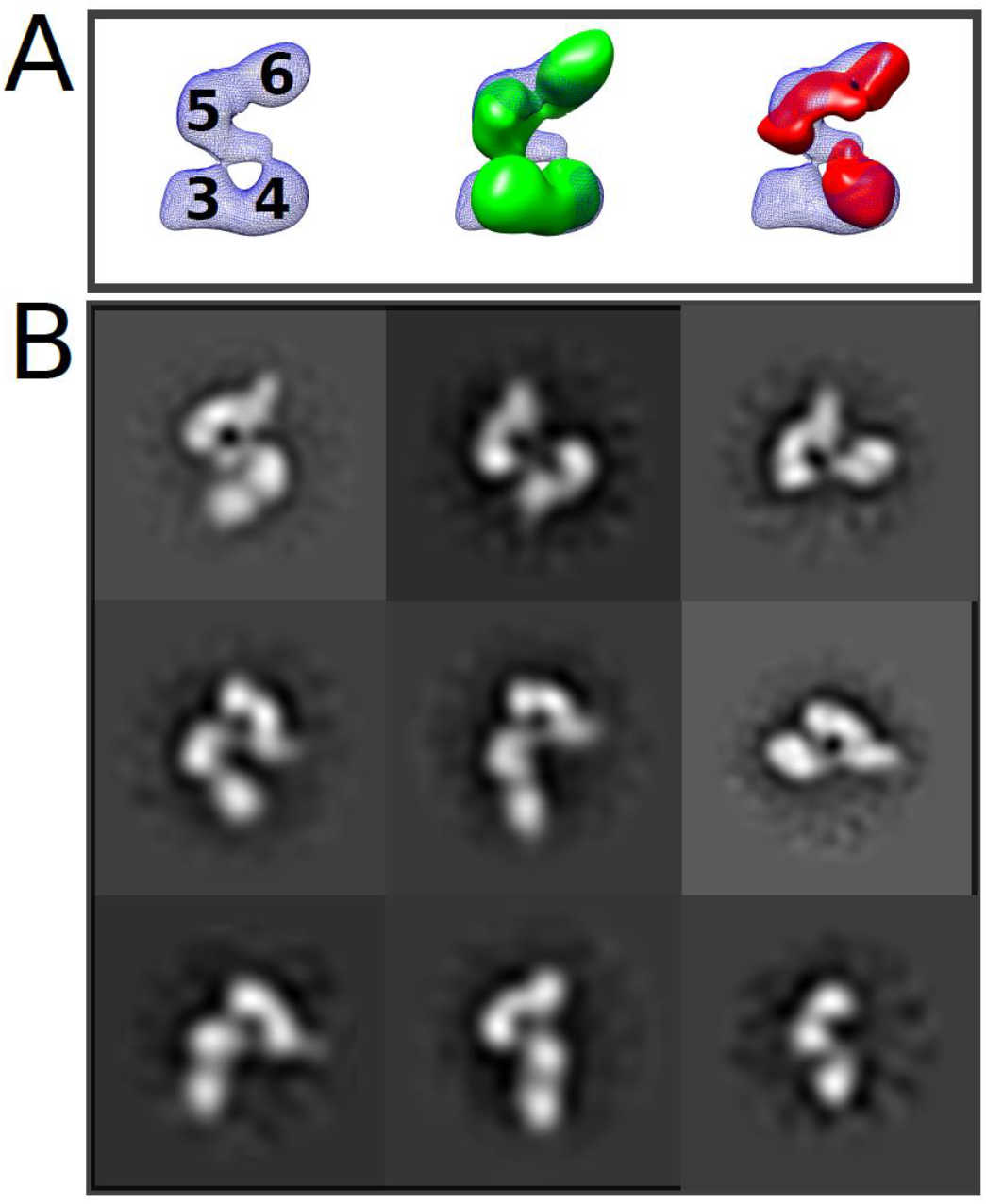
Superposition of the three N-terminal deletion constructs allows identification of DBL3x (3), DBL4ε (4), DBL5ε (5) and DBL6ε (6) to defined volumes in ID2b-DBL6ε. (A) DBL2b-DBL6ε (blue mesh; left panel) is used as the reference to place DBL3x-DBL6ε (green solid; middle panel) and DBL4ε-DBL6ε (red solid; right panel). (B) Placement of domains is consistent with features of 2D projections as featured in three equivalent 2D classes for DBL2b-DBL6ε (left column), DBL3x-DBL6ε (middle column) and DBL4ε-DBL6ε (right column).

**Figure 7.**
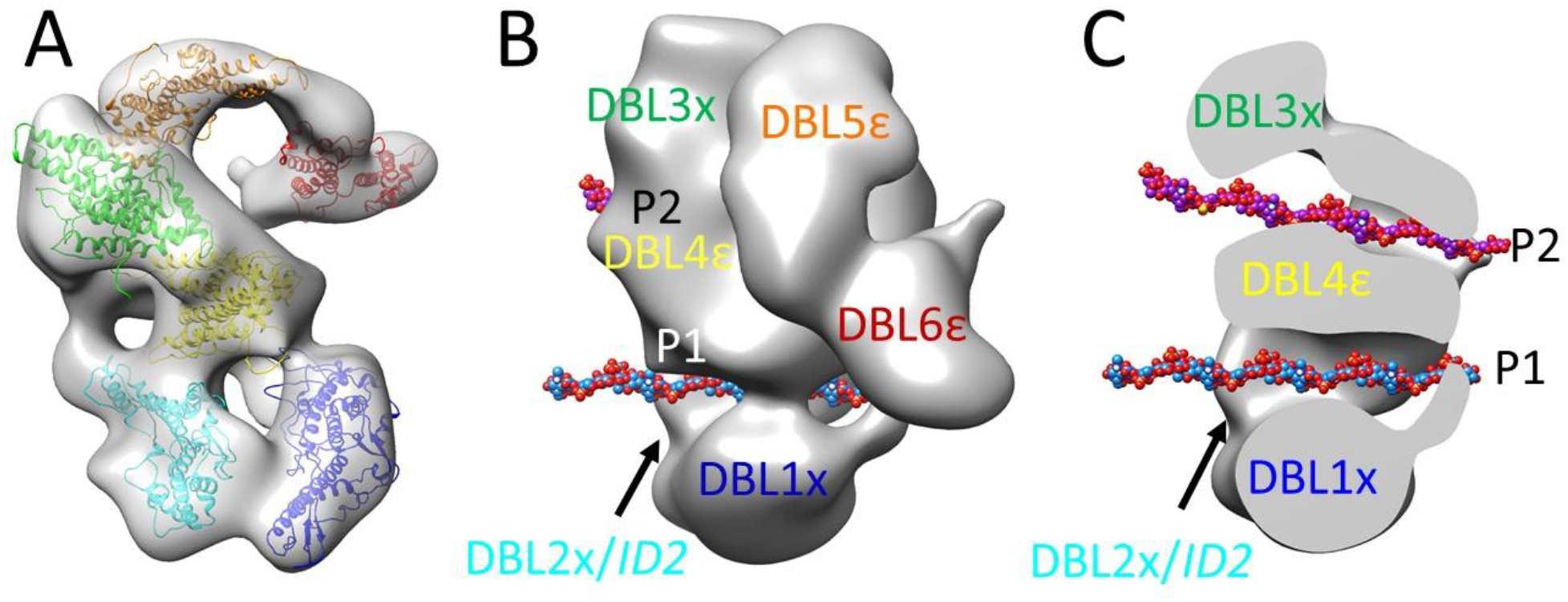
Placement of DBL1x-DBL6ε, based on the set of EM reconstructions, suggests a possible model for carbohydrate binding. (A) Transparent surface of the NTS-DBL6ε map (gray) into which DBL1x (blue), DBL2x (cyan), DBL3x (green), DBL4ε (yellow), DBL5ε (orange) and DBL6ε (red) have been positioned reveals that the 2 pores are formed by DBL1-DBL4ε. (B) Solid representation of NTS-DBL6ε map showing a 12mer CSA (PDB code 1CSA) in positioned in each pore, P1 and P2. (C) Transverse section through the NTS-DBL6ε with domains labeled as in (B).

The structural analysis of the EM and SAXS data were performed independently and thus a comparison of the models obtained from each technique provides reliable validation for the proposed structure. Adjusted for resolution, the volumes for each of the constructs generated by SAXS and negative stained EM are in good agreement. To further compare the models, EM density maps were converted to dummy atom models using EM2DAM (ATSAS 3.0) and theoretical SAXS curves were calculated from them and compared to the experimental SAXS curves (Fig. S10). There was good agreement between these curves and, as expected, the EM-based curve contained additional features consistent with finer detail and less movement or degeneracy.

### Docking of experimentally and computationally derived domain modules into the EM reconstructions

For rigid body fitting of data, models corresponding to DBL1x, DBL2x, DBL4ε and DBL5ε were generating using Chimera and Modeller (55,56). Structures of DBL3x and DBL6ε were used directly after modeling regions that were disordered in the crystal structures using the Chimera Modeller loops/refinement plug-in (56). However, ID1 (~200 amino acids) and ID2 (~300 amino acids) could not be modeled as suitable structural templates are not available (Table S3). The domains we placed as shown in Fig. 9A. DBL6ε fit into the expected density at the tip of the molecule and DBL5ε fit into the remaining density in the head. Based on the connection determined in the negative strain reconstruction,

DBL3x fit into the tail and DBL4ε was constrained to be adjacent to DBL3x and DBL5ε. DBL1x and DBL2x fit into the base, although the arrangement of these two domains is speculative because the folds of ID1 and ID2 are not known. DBL3x and DBL4ε can be accommodated in an arrangement similar to the structure of the DBL3x-DBL4ε tandem domains (44). Fitted in this way, the two large pores, P1 and P2, observed in NTS-DBL6ε, are created by DBL1x, ID1, DBL2x and DBL4ε, and DBL2x, DBL3x and DBL4ε, respectively.

## DISCUSSION

The VAR2CSA ectodomain and N- and C-terminal deletion constructs described in this work are produced in a mammalian expression system that exports them into the media as folded proteins. The purified proteins are very stable, as expected due to the presence of a large number of intra-domain disulfide bonds, and remain soluble and monomeric at concentrations >5 mg/ml, demonstrating that they are suitable for structural characterization. Analysis by SAXS and negative stain EM show that the protein constructs are homogeneous in terms of overall shape and apparent sizes. Taken together, this argues that the structures discussed here largely represent authentically folded VAR2CSA and folded domains thereof. Additionally, The *ab initio* envelope of NTS-DBL6ε calculated from SEC-SAXS, described here, contains the same features as models published earlier (38) after making allowances for the presence of the estimated 5% glycosylation (39) The *ab initio* envelope of NTS-DBL6ε calculated from SEC-SAXS, described here, contains the same features as published models after making allowances for the presence of the estimated 5% glycosylation (38,39). This suggests that slight differences in constructs and data collection/processing do not affect the overall structure of the molecule.

The single particle EM reconstructions of the ectodomain of VAR2CSA define the overall duck-like architecture of the molecule. Comparison of the SEC-SAXS bead models and negative stained EM reconstructions of the intact ectodomain and their various deletion constructs clearly define the locations of the DBL3x through DBL6ε domains and reveal the location of the DBL1x-ID2a module within this structure both in solution and on grids. The observed connectivity and overall structural details observed in the EM and SAXS data presented here reveals a packing of three tandem domains: DBL1x with DBL2x, DBL3x with DBL4ε and DBL5ε with DBL6ε. This packing is distinctly different from the SAXS-based model proposed by Clausen *et al* (39) that comprised of three tightly packed domain pairs DBL1x with DBL6ε, DBL2x with DBL5ε and DBL3x with DBL4ε and result in the N- and C-terminal domains co-localizing.

The new model of the ectodomain, described here, provides additional insight into how VAR2CSA might function in the context of infected red blood cells binding to C4S. Previous studies have shown that, in the knob-like structures observed on the surface of infected cells, VAR2CSA molecules are present largely at the tips (57,58). The identification of the C-terminus of the ectodomain with the anti-cMyc monoclonal antibody allows the orientation of the DBL domains to be determined with respect to the IRBC membrane (Fig. 6); an arrangement that places the head closest to the erythrocyte membrane and the body, and the feet of the ectodomain at the membrane-distal tip of the structure extending out from the knob into solution. This model is consistent with experimental data demonstrating that regions within NTS-DBLx-ID1-DBL2x and some portion of ID2, which comprise the feet, are required for specific binding of the C4S ligand. The proposed arrangement promotes relatively free access of the putative ligand-binding region to the receptor C4S chains attached to membrane bound portion and extracellular matrix of the placental intervillous blood space. By contrast, if DBL1x and DBL6ε were adjacent, as suggested in the previous model for domain organization (59), then the putative C4S-binding domains (DBL1x-DBL2x) would be located nearer to the knob membrane, making it less accessible to placental CSA chains.

Structurally, both the SAXS and negative stain reconstructions suggest that the intact ectodomain, isolated DBL1x-ID2a and, to a somewhat lesser extent, DBL3x-DBL6ε and DBL4ε-DBL6ε are structurally rigid at this resolution despite the paucity of obvious contacts between DBL5ε-DBL6ε and the body and tail of the structures (DBL3x, DBL4ε and some portion of ID2). Examination of the disulfide bonding pattern observed in the previously determined crystal structures identifies 8 canonical intra-domain disulfide bonds, although only one set (C(8)-C(12)) is observed in all the crystal structures of DBL3x, DBL4ε and DBL6ε. However, the remaining free, surface exposed, cysteine residues potentially are available to form inter-domain disulfide bonds that could stabilize the overall architecture of VAR2CSA. Although the current studies, due to their limited resolution, do not provide direct evidence for the existence of inter-domain disulfide bonds, at least two discrete volumes of stain-excluding density between the spatially distinct regions of the head and DBL4ε are observed in 2D classes of all DBL4ε-containing reconstructions. This is consistent with the presence of inter disulfide bonds and could account, in part, for the observed rigidity of this region of the structure. It should be noted that only one of these densities, which we term the neck, connecting one end of the head to DBL4ε is routinely observed in the 3D maps at the typical contour levels used. This most likely reflects a combination of conformational heterogeneity between the head and the body and limitations in image quality and number for the minor tilted views compared to the dominant side-on views.

A striking characteristic of the NTS-DBL6ε structure is the presence of two pores, P1 and P2 that are approximately parallel and traverse the width of the protein, accounting for the layers seen in some 2D classes. Both pores have dimensions that could accommodate CSA 10-12mers or other extended carbohydrate polymers (Fig. 9 B, C). In this speculative model, binding of carbohydrate in P1 would involve residues from DBL1x, ID1, DBL2x, DBL4ε and likely ID2a, and in P2 involve residues from DBL2x, DBL3x, DBL4ε and possibly ID2b. Enveloping the carbohydrate in a pore provides a larger surface area for binding interactions while minimizing the accessibility of this functionally-essential surface to immune surveillance. CSA and its non-sulfated homolog chondroitin can be described as stiff worm like polymers (persistence lengths ~70-80 Å) (60). Based on the structure of CSA (1CSA.PDB), a fully extended 12-mer CSA chain has an end-to-end distance of ~ 100-115 Å with an estimated minimum RMS end-to-end distance of ~90-100 Å, although it is likely to be closer to the fully extended length in solution. Therefore this single 12mer chain cannot occupy both pores simultaneously without either large scale rearrangements in the highly disulfide bond stabilized protein or substantial protein-induced hairpin bending of the CSA; either scenario seems unlikely. Many studies have concluded that residues within DBL1x-2IDa are necessary and sufficient for high affinity CSA binding, and the arrangement of domains forming P1 is consistent with this data. Binding studies presented here also show that DBL1x-2IDa has a lower affinity for CSA than NTS-DBL6ε, consistent with involvement of interactions outside of this region, such as DBL4ε in our model. The role of P2 in carbohydrate binding, if any, is less clear. One possibility is that it could be involved in protein-protein interactions either in the knob or during transport from the parasite to the surface of the red blood cell. A second possibility is that it is a carbohydrate-binding site, although to date, binding studies have not revealed the presence of two high affinity CSA binding sites. Thus, if each binds a distinct carbohydrate chain, then the second site would have to be of much lower affinity than the first and binding would have to be independent. Biologically, this might have some advantage for parasite attachment to the placental extracellular matrix. The CSPG molecules are part of a complex mixture of polymers that form the host extracellular matrix of the intervillous spaces. In the context of this complex host receptor environment, two binding sites on VAR2CSA may facilitate its attachment to a heterogeneous carbohydrate-dense environment. However, at the current resolution it is impossible to clearly determine which of these possibilities, if any, are correct.

In conclusion, these studies have revealed for the first time, the structure of VAR2CSA, at moderate resolution, its orientation relative to the IRBC membrane and the relative arrangement of domains. The structure contains two clear pores that transverse the molecule and suggest a possible model for host receptor binding. This model provides plausible explanations for the high affinity CSA interaction and the ability to evade initial immune surveillance.

### Experimental Procedures

#### Synthesis and sub-cloning of VAR2CSA gene into pSectTag2-based expression vectors

The sequence of the codon optimized VAR2CSA gene from strain 3D7 (JQ247428.1) was used as a starting point to generate a synthetic gene (amino acids 59-2641). Serine or threonine residues at 20 sites, identified for sites of potential *O*-linked glycosylation arising from the mammalian machinery (38), were replaced with codons for alanine or leucine (GeneArt, ThermoFisher Scientific). KpnI and ApaI sites were added to the 5’ and 3’ end, respectively for subcloning into the pSecTag2/Hygro2 vector which contains a N-terminal IgK secretion signal and non-cleavable C-terminal cMyc and hexahistidine tags for protein purification. In some constructs, as detailed in the Fig. 1A legend, the cMyc tag in the vector was replaced by a TEV-cleavable 3X-FLAG-tag using a GBLOCK primer listed in Table S1. In order to generate NTS-DBL6ε, a GBLOCK (IDT) (Table S1) corresponding to amino acids 1-58 was designed and inserted into pSecTag2/Hygro A vector using In Fusion (Takara, Japan) according to the manufacturer’s protocol to create a vector that contained amino acids 1-58 followed by KpnI and ApaI sites. Residues 59-2641 were then sub-cloned as before, which generated insertion of a glycine and threonine residue between native residues 58 and 59. DBL1x-ID2a (aa 59-1019), ID2b-DBL6ε (aa 1016-2641), DBL3x-DBL6ε (aa 1218-2630), DBL4ε-DBL6ε (aa 1560-2630) were created by PCR using the plasmid DNA of DBL1x-DBL6ε (1 ng/μl) as a template, primers (Table S1) and the Q5 site directed mutagenesis kit (NEB) for polymerase and parental DNA digestion. In all cases, after digestion 0.8 μl of the PCR reaction was introduced by transformation into 20 μl DH10B cells and grown at 37 °C on LB carbenecillin plates overnight. Single colonies were selected and inoculated into 8 ml LB media supplemented with carbenicillin (100 mg/ml). At mid log phase of growth, 1.5 ml was removed for freezer stocks, added to 150 μl autoclaved 80 % glycerol in 2 ml cryovials and stored at −80 °C. The remaining culture was grown for an additional ~2 h and harvested by centrifugation in a swinging bucket rotor for isolation of DNA using QIAprep Spin Miniprep kit (Qiagen), following the standard protocol. All DNA was sequenced through the coding region to confirm that the sequence was correct (Eurofins Genomics, Lancaster, PA). The translated protein sequences of the constructs are detailed in Table S2. Plasmid DNA was obtained by scraped from the −80 °C freezer stock into LB media supplemented with 100 mg/ml carbenicillin and isolated using the appropriate QIAprep kit.

#### Protein expression and purification

The expression and purification of proteins is based on Srivatsava *et al* (44) with minor modifications. Freestyle human embryonic kidney (HEK 293F) cells were grown in Freestyle 293 serum-free expression medium (Invitrogen Cat No. 12338-018) and the relevant expression plasmid was transfected using polyethylene amine (PEI), at a ratio of 1:3 DNA:PEI and grown at 37 °C. After 24 hours, cells were diluted with an equal volume of FreeStyle 293 medium, prewarmed to 37 °C and supplemented with valproic acid (Cat. No. P4543-100G, Sigma) to a final concentration of 2.2 mM. Protease inhibitors (Cat. No. K1008, APEX BioCat; final concentration 1x from a 200x stock) were added to the media after 48 h and the cultures were harvested 72 h post transfection, by centrifugation at 6000 × g, 4 °C. The clarified media, which contained the exported VAR2CSA proteins, was filtered through a 0.22 μm filter, Protease Inhibitor Cocktail (200x in DMSO, Apex Bio, K1008) added to a final concentration of 1x and the media either frozen at −20 °C or used directly for protein purification. Proteins were purified from fresh or thawed media by concentration to 50 ml using a VivaFlow 50 crossflow cassette MWCO 100 k (Cat. No. VF05C4, Sartorius), diluted with 20 mM imidazole + Ni Buffer A (500 mM NaCl, 10 mM NaKHPO_4_, pH 6.8), then reconcentrated to a final volume of 50 ml. The expressed proteins were isolated via a three-step procedure. Processed media was applied to a HisTrap Fast Flow HPLC column (Cat. No. 17531901, GE Healthcare), pre-equilibrated with Ni Buffer A + 20 mM imidazole and washed with 5 column volumes of this buffer + 35 mM Imidazole and the bound protein was eluted in a step gradient with Ni Buffer A + 300 mM Imidazole. The peak fractions containing VAR2CSA were concentrated using a Centricon 100 kDa and exchanged to 50 mM NaCl +10 mM NaKHPO_4_, pH 6.8 (CPX Buffer A,) prior to ion-exchange chromatography. The concentrated protein was loaded on an Eshmuno CPX column (8 × 20 mm, Sigma) pre-equilibrated with 50 mM NaCl+CPX Buffer A. The column was washed with 5 column volumes of 95% 50 mM NaCl+CPX Buffer A: 5% (1 M NaCl+CPX Buffer A) CPX Buffer B and the bound protein was eluted in a 10-column volume linear gradient from 40-80 % CPX Buffer B. The peak fractions were analyzed by SDS-PAGE and Coomassie staining. Protein-containing fractions were pooled and concentrated (Microsep, 100 kDa cutoff) to ~5 mg/ml and applied to a TSK-G3000SWxL column (7.8 × 30 cm, Tosoh Bioscience, King of Prussia, PA) pre-equilibrated with SEC Buffer A (10 mM NaKHPO_4_, pH 6.8, 150 mM NaCl). The protein was eluted with the same buffer and protein-containing fractions were analyzed by SDS-PAGE and Coomassie and/or silver staining. Fractions containing purified proteins (>95% pure) were concentrated to 2-8 mg/ml. The proteins were either used for experiments immediately or snap frozen in liquid nitrogen and stored at −80 °C for future use.

#### CD measurements

CD spectra (190-260 nm) of proteins in 0.2 mg/ml of 10 mM NaKHPO_4_, pH 6.8, 100 mM NaF were recorded on a Jasco J-1500 spectrophotometer at 25 °C and 90 °C using a 0.1 cm path length cuvette. Spectra were accumulated at 0.5 nm data intervals at a scan rate of 50 nm/min. For each sample, three scans were collected, averaged and corrected for base line rotation. Data were converted to molar ellipticity. For thermal denaturation, spectra of proteins in 0.2 mg/ml of 10 mM NaKHPO_4_, pH 6.8, 100 mM NaCl were recorded on a Jasco J-720 spectrophotometer spectra at 222 nm over the temperature range 25 °C to 85 °C at 1 °C intervals. Data was processed using GraphPad Prism version 5.

#### Analysis of VAR2CSA proteins binding to CSPG

The extent of binding of the VAR2CSA proteins was assessed in a sandwich ELISA-based assay using bovine corneal CSPG-II (49). The preparation of bovine corneal CSPG-II used here contains low-sulfated CSA chains with 28% 4-sulfated, 3 % 6-sulfated and 69 % non-sulfated disaccharide moieties. Briefly, specific wells of a 96-well microtiter plate were coated with bovine corneal CSPG-II by incubation with 50 μl (200 ng/ml CSPG-II in PBS, pH 7.2) at 4 °C overnight. All subsequent steps of this assay were performed at room temperature. Unbound CSPG-II was removed by washing the wells with 100 μl PBS, pH 7.2 containing 0.05% Tween-20 (PBST) and then all wells were blocked with 100 μl of 1% BSA in PBS, pH 7.2 for 2 h to minimize non-specific binding. Binding of the individual VAR2CSA proteins was assessed by serial 2-fold dilutions (50 μl) of the specified VAR2CSA construct (see Fig. S2 legend) into CSPG-II coated and uncoated (blank, to assess non-specific protein binding) wells of the microtiter plate and incubated for 2 h. Unbound protein was removed by washing of each well 3 times with 100 μl of PBST. Mouse anti-cMyc monoclonal antibody (Cat No. NB600-302, NOVUS Biologicals, Centennial, CO; cMyc mAb) was then added to each well (50 μl of 1:1000 diluted), washed three times with 100 μl PBST, and incubated with 50 μl of 1:10000 diluted HRP-conjugated goat anti-mouse IgG (Johnson Immunoresearch, West Grove PA) and then washed three times with 100 μl of PBST. The bound cMyc mAb was quantitated by the addition of 50 μl of 3, 3’5, 5’-tetramethylbenzidine substrate to all wells and the color allowed to develop for 20 min at room temperature. The incubation time was chosen to ensure that the assays were in the linear detection range. Color development was stopped by addition of 25 μl of 2.5 M HCl. The extent of protein bound to the CSPG-coated wells was estimated from the absorbance at 450nm.

#### Inhibition of VAR2CSA proteins binding to CSPG by various glycosaminoglycans (GAGs)

A modified version of the assay outlined above was used to analyze the ability of GAGs to inhibit binding of NTS-DBL6ε and DBL1x-ID2a binding to CSPG-II. The following GAGs were evaluated: bovine tracheal CSA (54 % 4-sulfated, 39 % 6-sulfated, 8 % non-sulfated; Cat. No. C-8529, Sigma), C6S (~20 % 4-sulfated, ~80 % 6-sulfated disaccharide moieties, Seikagaku Corp, Japan) and HA (100 % non-sulfated Cat. No. H-1504, Sigma). Microtiter plates with bovine corneal CSPG-II, prepared as described above, were used. Separately, NTS-DBL6ε and DBL1x-ID2a protein (2 μg/ml in PBS, pH 7.2) was incubated at room temperature with GAGs at concentrations ranging from 0.03 μg/ml to 5 μg/ml. After 40 min, 50μl of each solution was added to the CSPG-coated plates and incubated for 1 h. The extent of protein binding to the CSPG-coated wells was measured as described above.

#### Small angle X-ray scattering measurements

Data were collected by in-line SEC-SAXS either at LiX 16-ID beamline of the National Synchrotron Light Source II (NSLSII), Brookhaven National Laboratory, Upton, NY 11973 or 18ID of the Advanced Photon Source (APS), Argonne National Laboratory, Argonne, IL (Table 1, S3). For data collection, 250 μl of each sample was applied to a Superose 6 10/300 GL size exclusion column (GE healthcare) at concentrations between 4-10 mg/ml and eluted at room temperature with a flow rate of 0.5 ml/min in 10 mM NaKHPO_4_, pH 6.8, 150 mM NaCl. Scattering curves were collected continuously throughout the elution to obtain buffer and protein scattering curves. Data were initially processed using beamline software py4XS (NSLSII) and RAW (APS) and subsequently programs in the ATSAS Suite (versions 2.8 and 3.0) (53). *Ab initio* bead models were calculated from 20 candidate models calculated in DAMMIF, averaged using DAMAVER and refined with DAMMIN. For display purposes and superposition, the bead models were converted to volumes within Chimera and superimposed using the fitmap algorithm (55,56,61)

#### Preparation of NTS-DBL6ε:anti-cMyc antibody complex and DBL1x-ID2a:anti-cMyc antibody complex for EM

cMyc mAb (Cat NB600-302, Novus Biologicals, CO) and NTS-DBL6ε or DBL1x-ID2a proteins were mixed at a ratio of 1:2 and incubated for 30 min on ice. Solutions were loaded onto TGX3000S size-exclusion column (Tosho Biosciences) and eluted with 10 mM NaKHPO_4_, pH 6.8, 150 mM NaCl at a flow rate of 0.5 ml/min. Fractions containing protein:antibody complexes were identified by the absorbance profile and SDS-PAGE of individual fractions, pooled, concentrated to 0.5 mg/ml and used for negative staining for EM.

#### Negative staining and EM analysis

Formvar-coated carbon grids (300 μm mesh; EMS, PA) were prepared by plasma cleaning (air) for 60 s using an Emitech K590X plasma cleaner (Diener Electronics, Germany) immediately prior to use. For NTS-DBL6ε, or DBL1x-2IDa, a 4 μl aliquot of the protein alone, or bound to cMyc mAb (25 μg/ml in 10 mM NaKHPO_4_, pH 6.8, 150 mM NaCl) where relevant, was applied to the grids and incubated at room temperature for 10 s. Excess protein solution was removed from the edge of the grid using thin strips of Whatman No. 1 paper. The grids were washed 3 times by touching them sequentially to the surface of three distilled water drops (35 μl) and excess liquid removed as before. For staining, the grids were floated sequentially on the top of three 35 μl drops of 1% uranyl acetate in water (35 μl) for 2x 10 s, and 1 min, respectively, air dried at room temperature for 1 h and stored under anhydrous conditions until needed. EM data were collected using a JEOL 2100 transmission electron microscope equipped with a 4K CCD camera. All images were collected at 200 eV at a pixel size of 2.3 Å/pixel using Digital Micrograph.

#### Single particle reconstruction from EM micrographs of negative stain images

Single particle reconstructions were performed in Relion 3.1 (62) and validated using tools and protocols in Scipion 2 (63). Particles from 5-10 images were picked automatically using a range of minimum and maximum particle diameters and thresholds with the Lorenztian of Gaussian algorithm to estimate optimal values. Particles were then picked from the entire set of images using the same algorithm and the optimized numbers for each parameter and extracted, using a box size of 1.5 times the longest dimension of the particle, with a pixel size of 6.9 Å to improve signal to noise. The particle stack was subjected to 2-3 rounds of 2D classification to obtain a starting particle set for 3D reconstruction. Initial starting models were computed to 25 Å resolution using a randomly chosen subset of 4000 particles. Subsequently, all of the particles were used in 3D classification with the calculated starting model as a reference. The best class or classes, were subjected to 3D refinement and post processed to obtain a final masked map. The reported resolution was obtained using the gold standard FSC (gsFSC) during post processing. In some cases, a second round of 2D/3D classification and refinement improved the shape of the gsFSC. In these cases, the particles from the first 3D refinement were subjected to an additional round of 2D classification followed by 3D classification with a single class. The resultant map was further refined and post-processed, as before, to yield the final map described in this work. The maps for all constructs were validated using the 3D reprojection and overfitting tools in Scipion 2 with data from the final refinement in Relion 3.1 to confirm that there was no overfitting within the resolution ranges used. A summary of the results of reconstruction and 3D reprojection are given in Table 2 and Fig. S4-9.

#### Basic homology modeling of individual domains

Templates for homology modeling were identified using the BLAST server (https://blast.ncbi.nlm.nih.gov/Blast.cgi) and visualized using Chimera. Basic homology modeling was performed using the MODELLER interface within Chimera. For each domain, models were evaluated by visual inspection, and GA341 and zDOPE scores (Table S4). Residues 1-9, 402-568, 906-1204, 1945-1978 and 2284-2395 did not have credible templates and are not included in the final assembly.

## Acknowledgements

We thank Dr Mark Yeager, University of Virginia for hosting a sabbatical study (JMF) and Drs. Michael Purdy, Ali Khan, Susan Leonhardt, Maciej Jagielnicki, Kelly Dryden and the rest of the members of the Yeager lab for their patience, kindness and friendship. We thank beamline staff at NSLS-II including Drs Vivian Stojanoff, Sirish Chodankar and James Byrnes for beamline support and Dr Lin Yang for thoughtful discussions and suggestions. We also thank Dr Suresh Nayaranan for help collecting SAXS data at the Beyond R_g_ BioSAXS short course at APS. Dr Matt Swullius is thanked for help with the EM studies.

## Statement of contributions

LG performed all protein purification and enzyme assays, performed and analyzed all CD experiments, wrote and edited the materials and methods for these sections and collected SAXS and EM data. MCB collected SAXS data, processed and evaluated all SAXS data and performed basic modeling, JMF processed and evaluated all EM data. MCB, CDG and JMF conceived the experiments, analyzed the results, formulated the discussion and wrote the manuscript. All authors have read the manuscript and agree with the conclusions presented.

## Conflict of Interest

The authors declare that they have no conflicts of interest with the contents of this article.

## FOOTNOTES

The studies performed were supported by grant R01 AI104844 (D.C.G.) from the National Institute of Allergy and Infectious Diseases of the NIH. This research used ID-16 of the National Synchrotron Light Source II, a U.S. Department of Energy (DOE) Office of Science User Facility operated for the DOE Office of Science by Brookhaven National Laboratory under Contract No. DE-SC0012704. The Life Science Biomedical Technology Research resource is primarily supported by the National Institute of Health (NIH), National Institute of General Medical Sciences (NIGMS) through a Biomedical Technology Research Resource P41 grant (P41GM111244), and by the DOE Office of Biological and Environmental Research (KP1605010). This research also used resources of the Advanced Photon Source, a U.S. Department of Energy (DOE) Office of Science User Facility operated for the DOE Office of Science by Argonne National Laboratory under Contract No. DE-AC02-06CH11357 and the beamline and its resources was supported by grant 9 P41 GM103622 from the NIGMS of the NIH. Molecular graphics and analyses performed with UCSF Chimera, developed by the Resource for Biocomputing, Visualization, and Informatics at the University of California, San Francisco, with support from NIH P41-GM103311.

## Notes

### Competing Interest Statement

The authors have declared no competing interest.

