## Supplemental Information for "Molecular architecture and domain arrangement of the placental malaria protein VAR2CSA suggests a model for receptor binding"

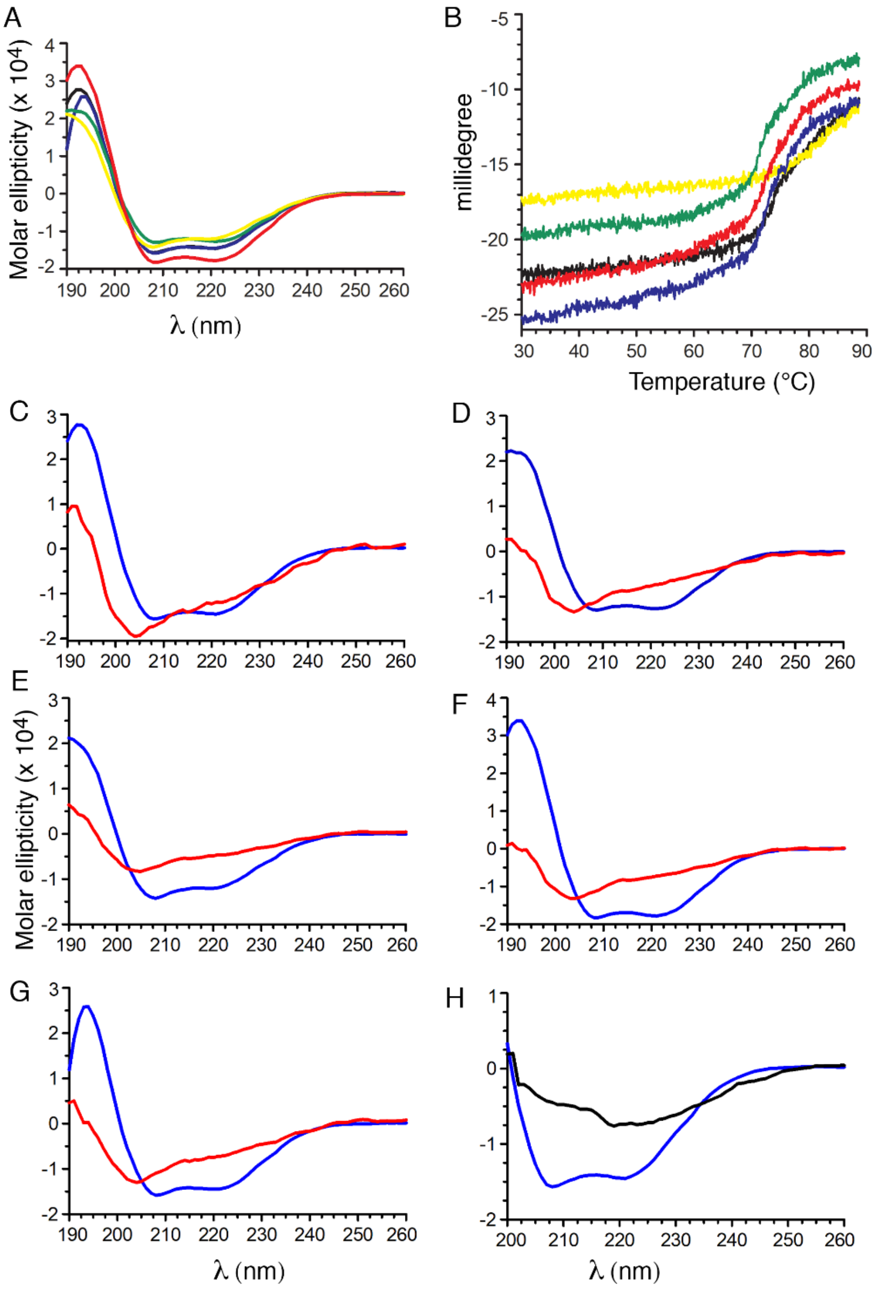


**Figure S1.** The recombinant VAR2CSA ectodomain and deletion constructs are folded and thermally stable. (A) CD spectra NTS-DBL6ε (black), DBL1x-ID2a (yellow), ID2b-DBL6ε (blue), DBL3x-DBL6ε (green), and DBL4ε-DBL6ε (red) are consistent with proteins containing α-helices. (B) Thermal unfolding curves of the constructs, colored as in (A). (C-G) CD spectra of proteins at 25 °C (blue trace) and 90 °C (red trace) for (C) NTS-DBL6ε, (D) DBL1x-2IDa, (E) ID2b-DBL6ε, (F) DBL3x-DBL6ε and (G) DBL4ε-DBL6ε. (H) NTS-DBL6ε at 25 °C (blue) and NTS-DBL6ε+TCEP at 25 °C (red).

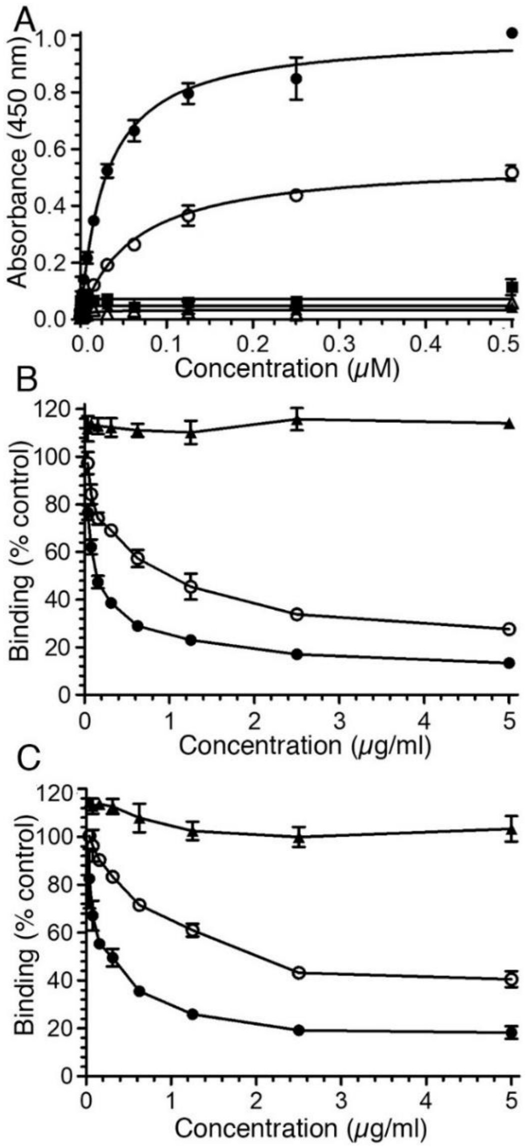


**Figure S2.** NTS-DBL6ε and DBL1x-2IDa bind C4S. (A) The binding of NTS-DBL6ε and deletion constructs, as described, to CSPG was assessed by an ELISA-based assay. The data were fitted to a non-linear regression 1 site specific binding in PRISM v 7.0. Data points correspond to the following constructs: (●) NTS-DBL6ε; (○) DBL1x-2IDa; (▲) ID2b-DBL6ε; (Δ) DBL3x-DBL6ε; (■) DBL4ε-DBL6ε. (B) Inhibition of NTS-DBL6ε and (C) DBL1x-2IDa binding to CSPG by glycosaminoglycans (●) CSA, (○) C6S and (▲) HA. Error bars represent SD from 3 technical replicates and is representative of three independent analyses.

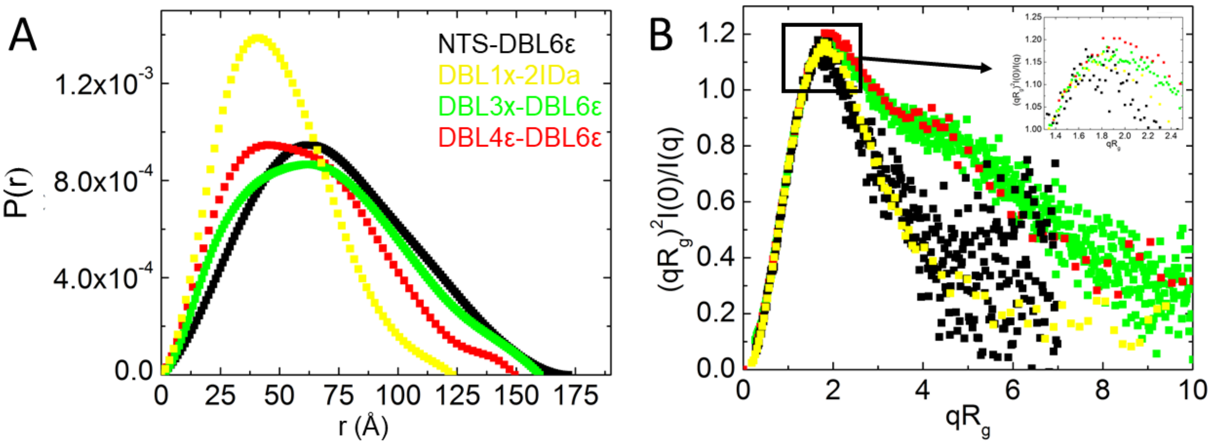


**Figure S3** Additional SAXS plots. (A) P(r) plot normalized to area for each of the protein constructs. (B) Dimensionless Kratky plot, colored as in (A) shows a peak at $qR_g$ ~1.7, as expected for folded proteins. DBL3x-DBL6ε and DBL4ε-DBL6ε show an additional shoulder at $R_g$, typical of protein containing distinct masses connected by a linker. Inset shows a more detailed view of the peak, covering the region shown in the black box.

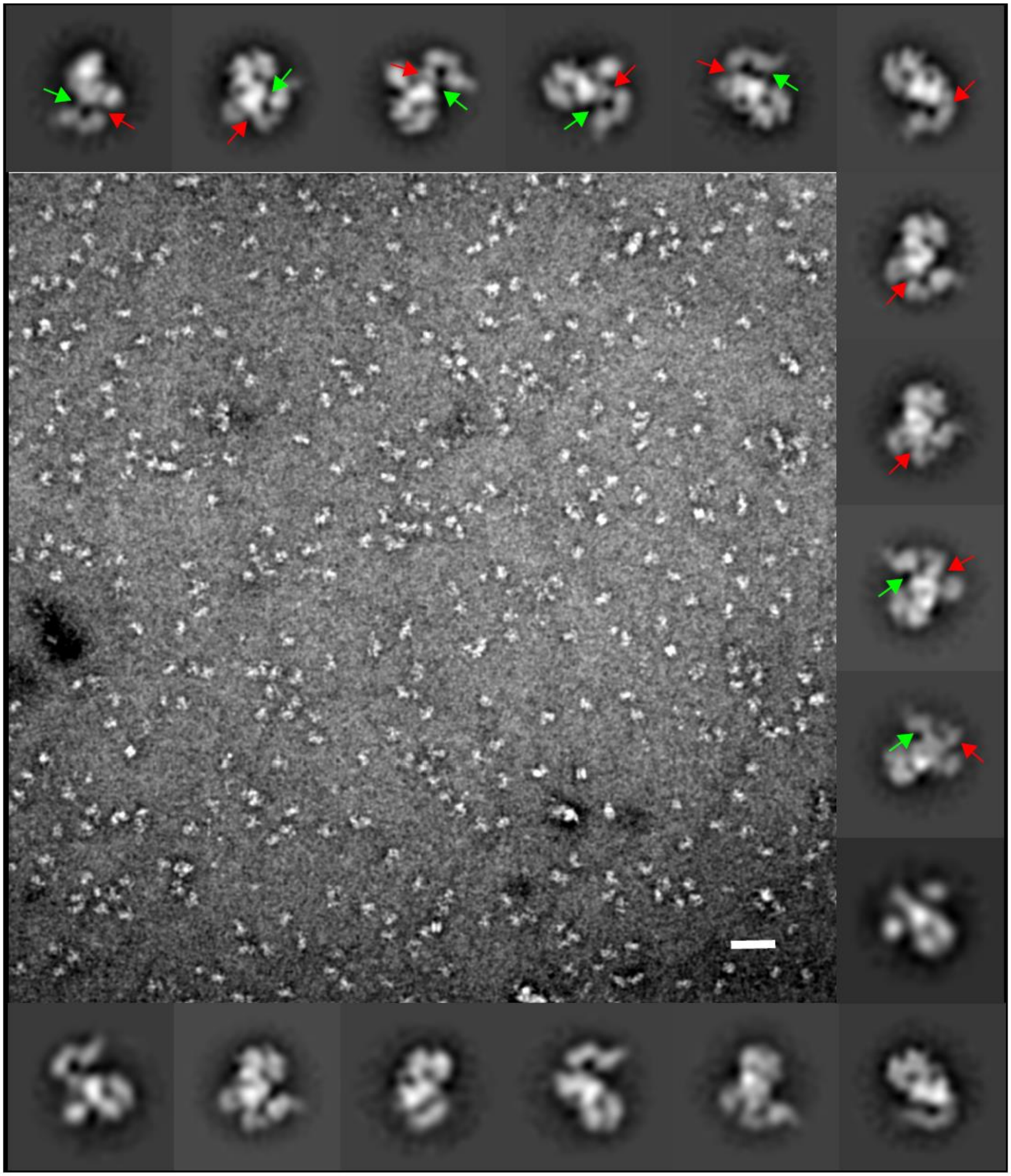

**Figure S4.** Micrograph and 2D classes of NTS-DBL6ε. (A) Sample micrograph containing particles of uranyl acetate stained NTS-DBL6ε. The white bar is 50 nm. A gallery of 2D classes obtained following the second round of 2D classification, surround the micrograph. Arrows denote the locations of the neck (red) and a second stain-excluding area (green) described in the text.

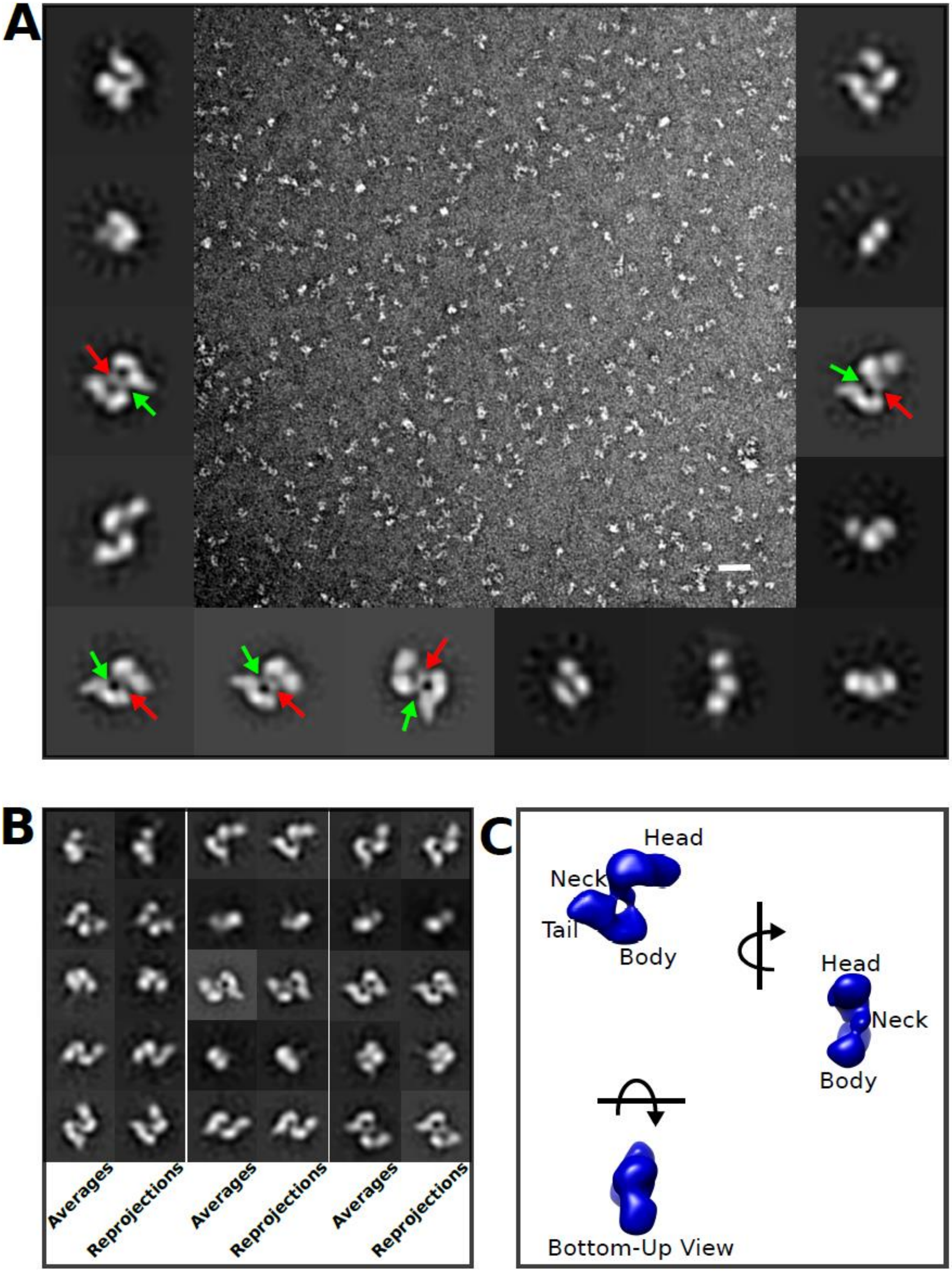
A
B
Averages
Reprojections
Averages
Reprojections
Averages
Reprojections
C
Head
Neck
Tail
Body
Head
Neck
Body
Bottom-Up View

**Figure S5**. Data and 3-D reconstruction of ID2b-DBL6ε. (A) Sample micrograph containing particles of uranyl acetate stained ID2b-DBL6ε. The white bar is 50 nm. A gallery of 2D classes obtained following the second round of 2D classification, surround the micrograph. (B) The top 15 2D-classes, as defined in Fig. 4, paired with the corresponding back projection of the 3D reconstruction. (C) Orthogonal views of the 3D reconstruction.

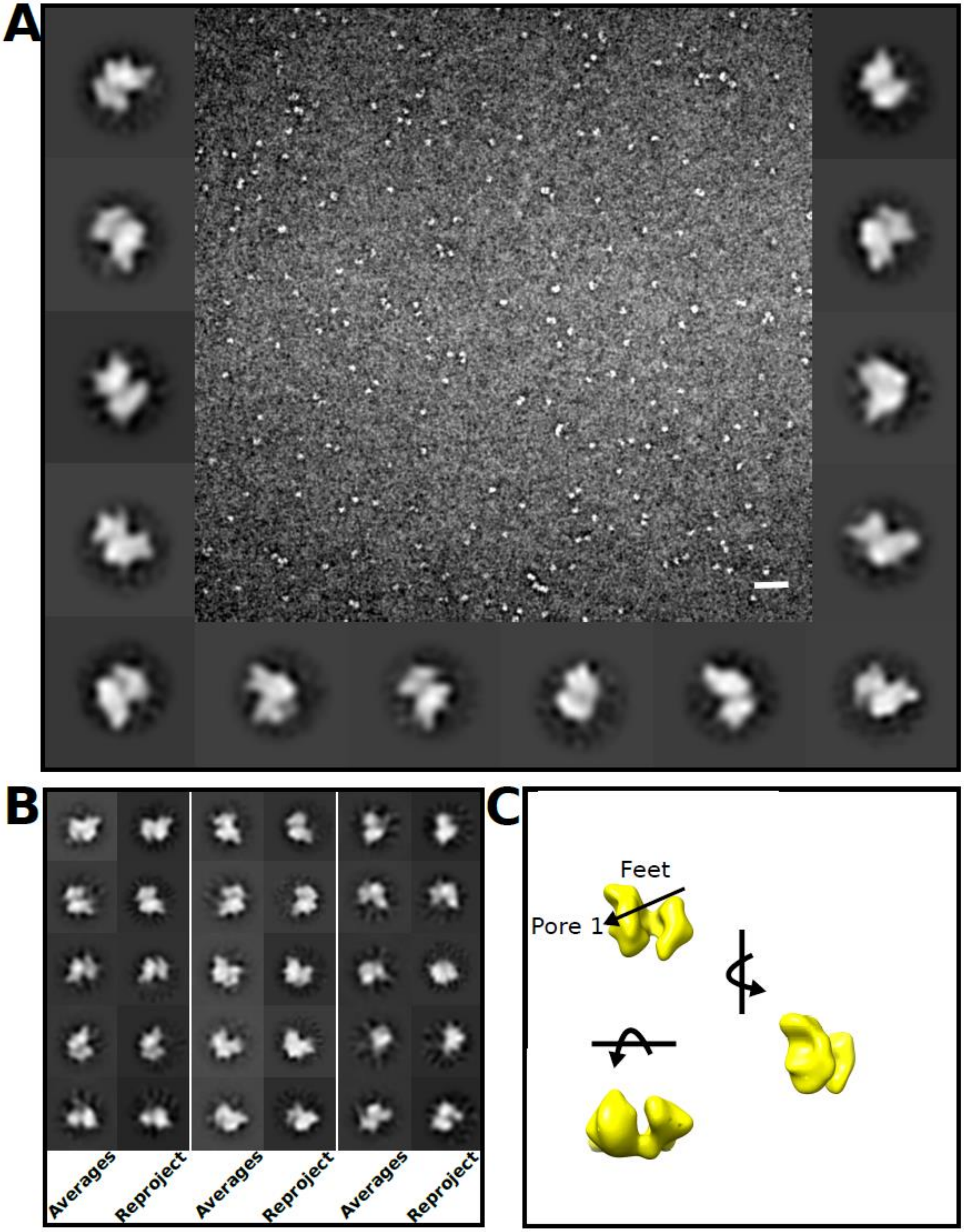


**Figure S6**. Data and 3-D reconstruction of DBL1x-2IDa. (A) Sample micrograph containing particles of uranyl acetate stained DBL1x-2IDa.  A galley of 2D classes surround the micrograph. (B) The top 25 2D-classes paired with the corresponding back projection of the 3D reconstruction. (C) Orthogonal views of the 3D reconstruction.

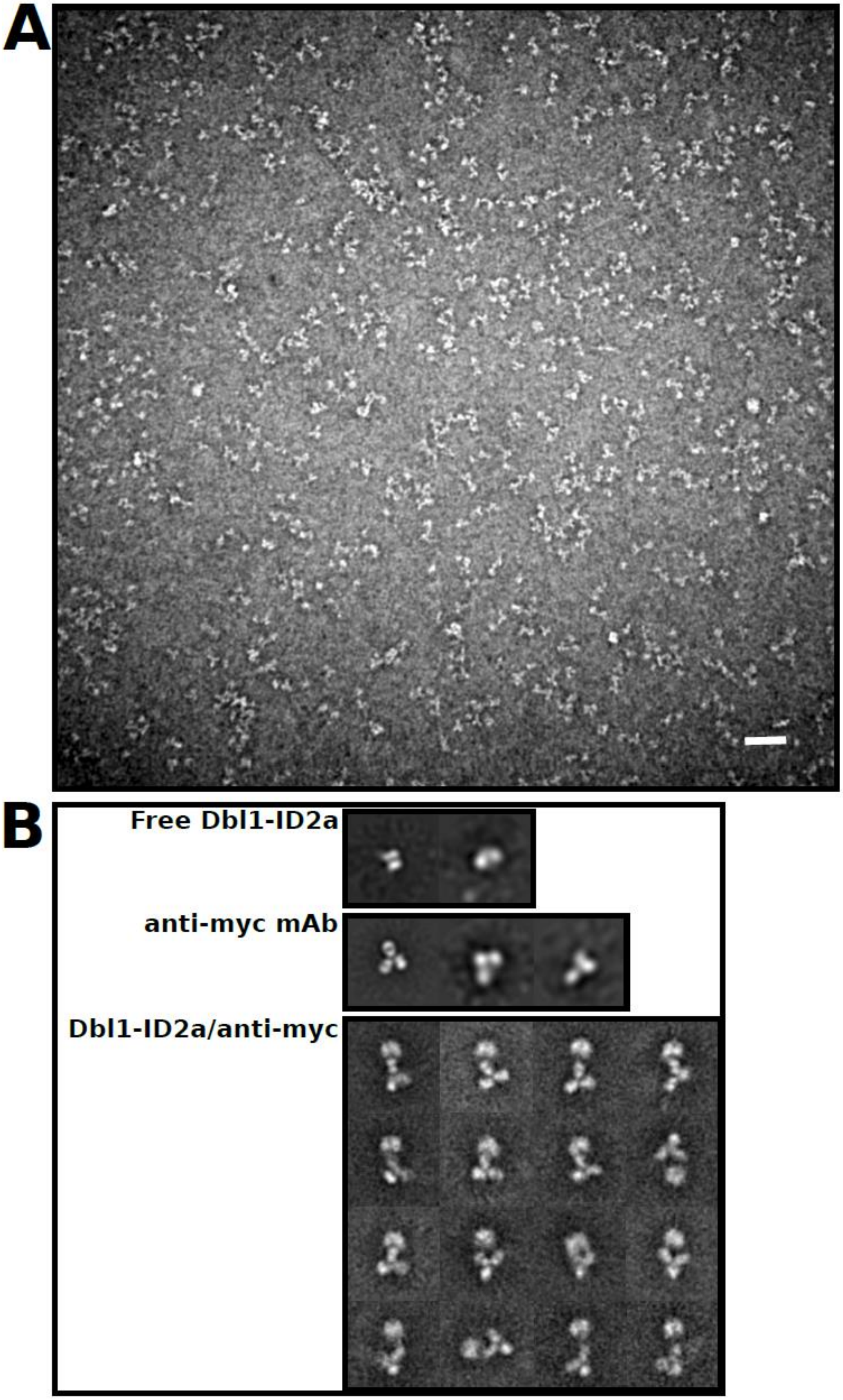
A
B
Free Dbl1-ID2a
anti-myc mAb
Dbl1-ID2a/anti-myc

F**igure S7**. Data and 3-D reconstruction of DBL1x-2IDa:anti cMyc antibody complex. (A) Sample micrograph containing particles of uranyl acetate stained DBL1x-2IDa:anti cMyc antibody. (B) The 2D-classes observed for DBL1x-ID2a alone, anti-cMyc antibody alone and the DBL1x-ID2a:anti-cMyc antibody complex. ). The location of the C-terminus could not be unambiguously assigned to any structural feature in this complex. This is likely due to the local conformational freedom of mAb in its interaction with the C-terminal tag of DBL1x-2IDa that are of nearly equal masses and the limited number of views of the complex that arise from preferred orientations in these images.

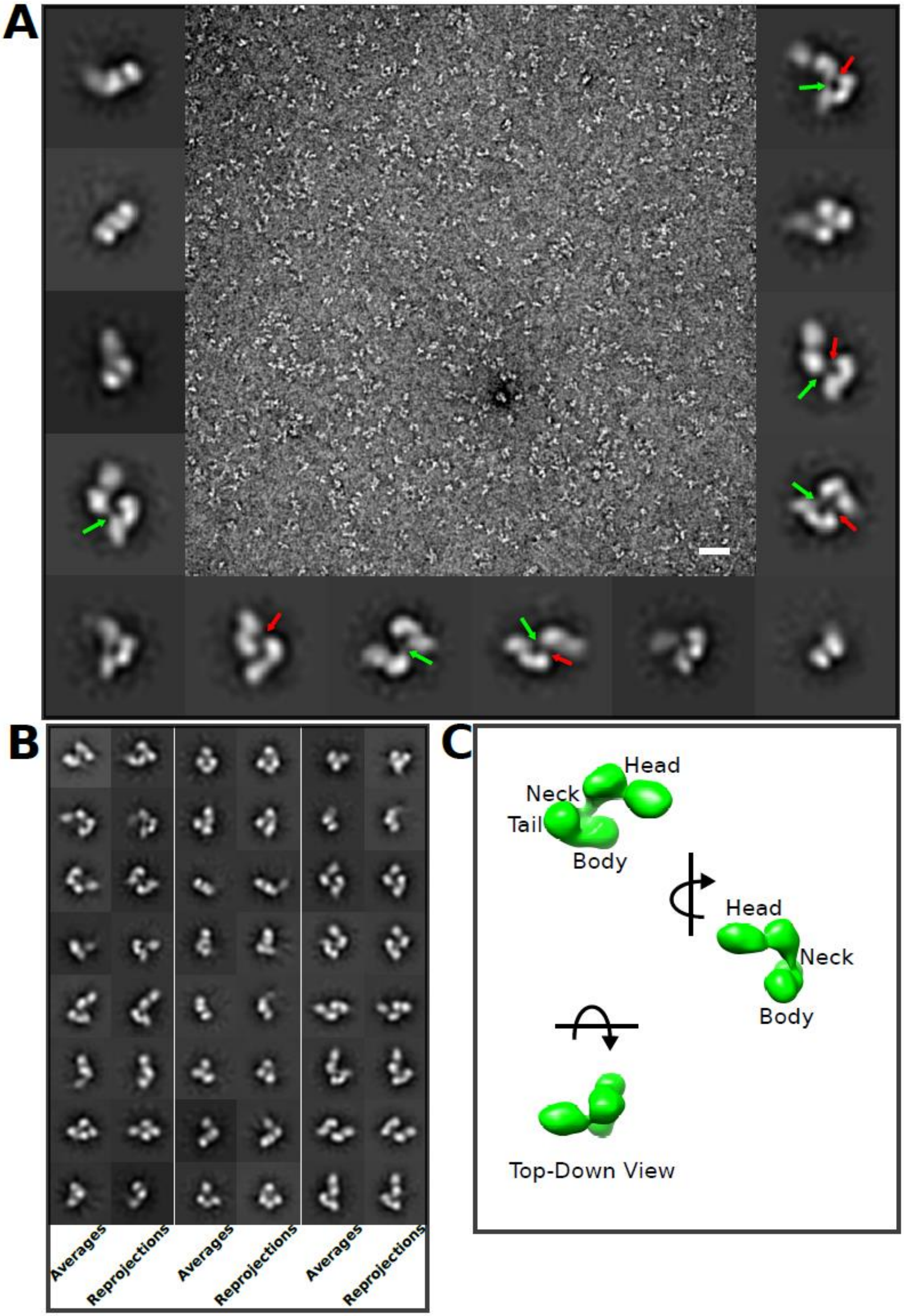
A
B
C
Averages
Reprojections
Averages
Reprojections
Averages
Reprojections
Head
Neck
Tail
Body
Head
Neck
Body
Top-Down View

**Figure S8.** Data and 3-D reconstruction of DBL3x-DBL6ε. (A) Sample micrograph containing particles of uranyl acetate stained DBL4ε -DBL6ε. A galley of 2D classes surround the micrograph. (B) The top 16 2D-classes paired with the corresponding back projection of the 3D reconstruction. (C) Orthogonal views of the 3D reconstruction.

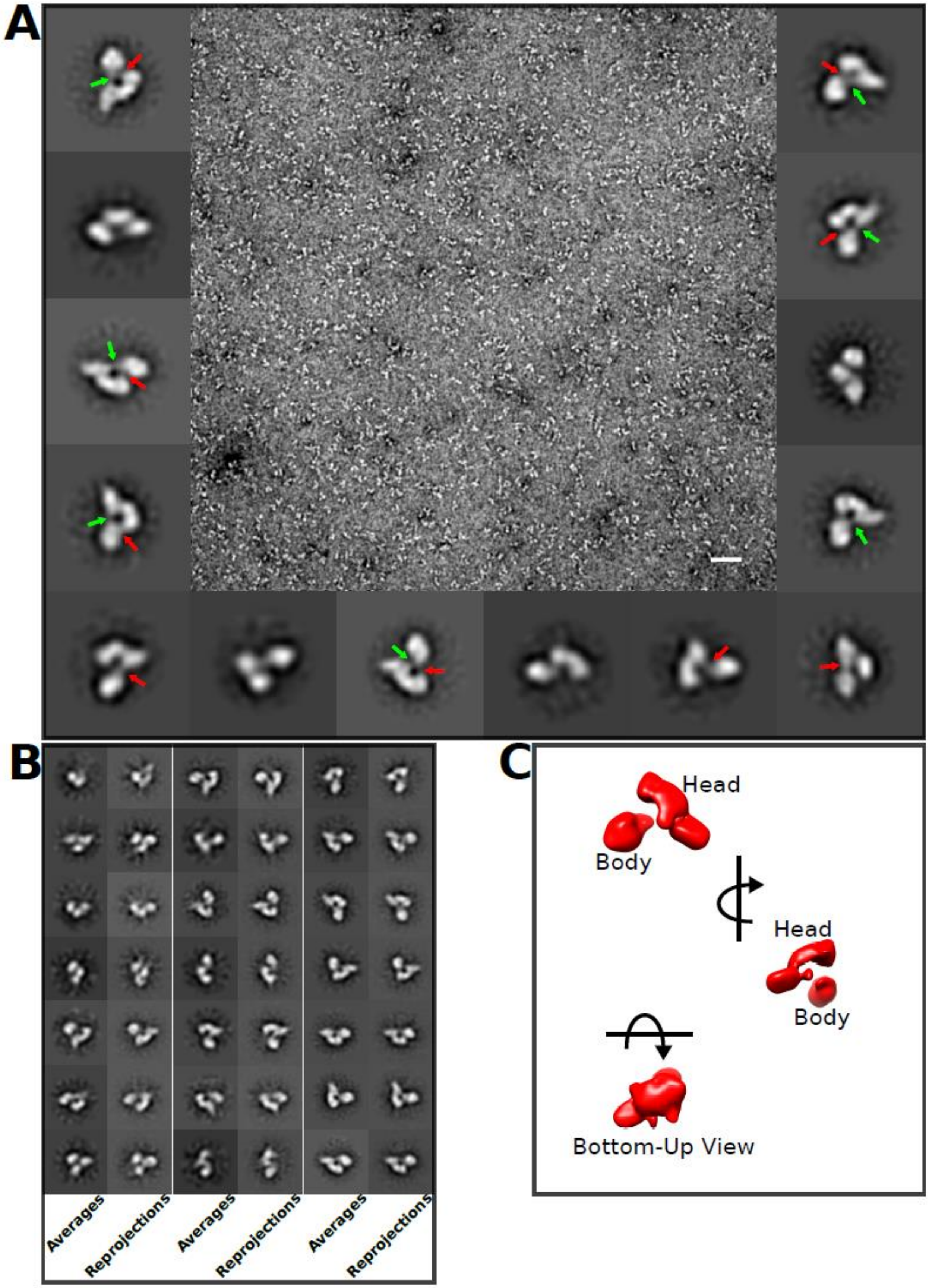
A
B
C
Averages
Reprojections
Averages
Reprojections
Averages
Reprojections
Head
Body
Head
Body
Bottom-Up View

**Figure S9.** Data and 3-D reconstruction of DBL4ε -DBL6ε. (A) Sample micrograph containing particles of uranyl acetate stained DBL4ε -DBL6ε.  A galley of 2D classes surround the micrograph. (B) The top 16 2D-classes paired with the corresponding back projection of the 3D reconstruction. (C) Orthogonal views of the 3D reconstruction.

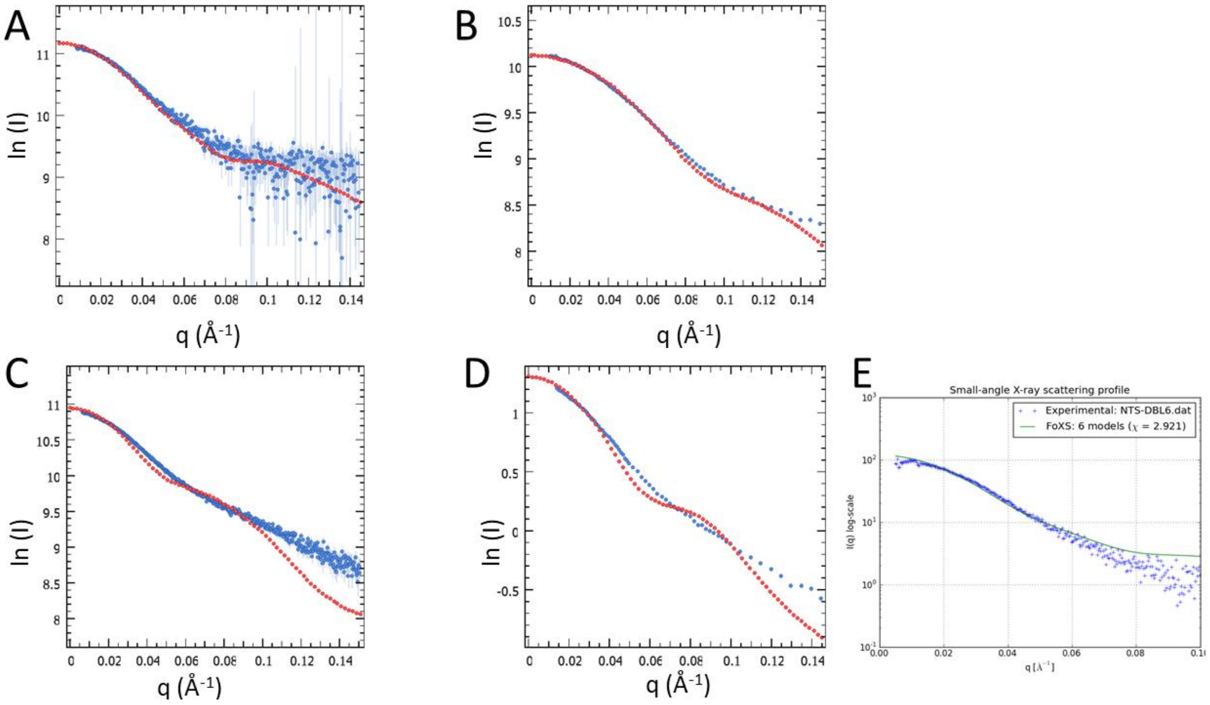


**Figure S10** Comparison of EM model based calculated scattering intensities (red) to experimental SAXS intensifies (blue) for the (A) NTS-DBL6ε (B) DBL1x-2IDa (C) DBL3x-DBL6ε (D) DBL4ε-DBL6ε constructs. Error bars corresponding to the SAXS data are drawn as blue vertical lines. (E) Fit of EM based homology model to experimental SAXS curve for NTS-DBL6ε.

**NTS-DBL6ε**

DKSSIANKIEAYLGAKSDDSKIDQSLKADPSEVQYYGSGGDGYYLRKNICKITVNHGTDAAQPA
RRARRTKLGTDSGTNDPCDRIPPPYGDNDQWKCAIILSKVSEKPENVFVPPRRQRMCINNLEKLN
VDKIRDKHAFLADVLLTARNEGERIVQNHPDTNSANVCNALERSFADIADIIRGTDLWKGTNSNL
EQNLKQMFAKIRENDKVLQDKYPKDQNYRKLREDWWNANRQKVWEVITCGARSNDLLIKRG
WRTSGKSNGDNKLELCRKCGHYEEKVPTKLDYVPQFLRWLTEWIEDFYREKQNLIDDMERHRE
ECTSEDHKSKEGTSYCSTCKDKCKKYCECVKKWKSEWENQKNKYTELYQQNKNEASQKNTSR
YDDYVKDFFKKLEANYASLENYIKGDPYFAEYATKLSFILNSADANNPSEKIQKNNDEVCNCNE
SGIASVEQEQISDPSSNKACITHSSIKANKKKVCKHVKLGVRENDKDLRVCVIEHTSLSGVENCCC
QDFLRILQENCADNKAGSSSNGACNNKNQEACEKNLEKVLASLTNCYKCDKCKSEQSKKNNKN
WIWKKSSGKEGGLQKEYANTIGLPPRTQSLCLVVCLDEKGKKTQELKNIRTNSELLKEWIIAAFH
EGKNLKPSHEKKNDDNGKKLCKALEYSFADYGDLIKGTSIWDNEYTKDLELNLQKIFGKLFRKY
IKKNNAAEQDTSYSSLDELRESWWNTNKKYIWLAMKHGAGMNSTTCCGDGSVTGSGSSCDDIP
TIDLIPQYLRFLQEWVEHFCKQRQEKVKPVIENCKSCKESGGTCNGECKTECKNKCEVYKKFIED
CKGGDGTAGSSWVKRWDQIYKRYSKYIEDAKRNRKAGTKNCGPSSTTNAAENKCVQSDIDSFF
KHLIDIGLTTPSSYLSIVLDDNICGADKAPWTTYTTYTTTEKCNKETDKSKLQQCNTAVVVNVPS
PLGNTPHGYKYACQCKIPTNEETCDDRKEYMNQWSCGSARTMKRGYKNDNYELCKYNGVDV
KPTTVRSNSAKLDDKDVTFFNLFEQWNKEIQYQIEQYMTNTKISCNNEKNVLSRVSDEAAQPKF
SDNERDRNSITHEDKNCKEKCKCYSLWIEKINDQWDKQKDNYNKFQRKQIYDANKGSQNKKV
VSLSNFLFFSCWEEYIQKYFNGDWSKIKNIGSDTFEFLIKKCGNDSGDGETIFSEKLNNAEKKCKE
NESTNNKMKSSETSCDCSEPIYIRGCQPKIYDGKIFPGKGGEKQWICKDTIIHGDTNGACIPPRTQN
LCVGELWDKRYGGRSNIKNDTKESLKQKIKNAIQKETELLYEYHDKGTAIISRNPMKGQKEKEE
KNNDSNGLPKGFCHAVQRSFIDYKNMILGTSVNIYEYIGKLQEDIKKIIEKGTTKQNGKTVGSGA
ENVNAWWKGIEGEMWDAVRCAITKINKKQKKNGTFSIDECGIFPPTGNDEDQSVSWFKEWSEQ
FCIERLQYEKNIRDACTNNGQGDKIQGDCKRKCEEYKKYISEKKQEWDKQKTKYENKYVGKSA
SDLLKENYPECISANFDFIFNDNIEYKTYYPYGDYSSICSCEQVKYYEYNNAEKKNNKLLCHEKG
NDRTWSKKYIKKLENGRTLEGVYVPPRRQQLCLYELFPIIIKNKNDITNAKKELLETLQIVAEREA
YYLWKQYHAHNDATYLAHKKACCAIRGSFYDLEDIIKGNDLVHDEYTKYIDSKLNEIFDSSNKN
DIETKRARTDWWENEAIAVPNIAGANKADPKTIRQLVWDAMQSGVRKAIDEEKEKKKPNENFPP
CMGVQHIGIAKPQFIRWLEEWTNEFCEKYTKYFEDMKSNCNLRKGADDCDDNSNIECKKACAN
YANWLNPKRIEWNGMSNYYNKIYRKSNKESEDGKDYSMIMEPTVIDYLNKRCNGEINGNYICCS
CKNIGENSASGTVNKKLQKKETQCEDNKGPLDLMNKVLNKMDPKYSEHKMKCTEVYLEHVEE
QLKEIDNAIKDYKLYPLDRCFDDKSKMKVCDLIGDAIGCKHKTKLDELDEWNDVDMRDPYNKY
KGVLIPPRRRQLCFSRIVRGPANLRNLKEFKEEILKGAQSEGKFLGNYYNEDKDKEKALEAMKNS
FYDYEYIIKGSDMLTNIQFKDIKRKLDRLLEKETNNAEKVDDWWETNKKSIWNAMLCGYKKSG
NKIIDPSWCTIPTTETPPQFLRWIKEWGTNVCIQKEEHKEYVKSKCSNVANLGAQESESKNCTSEI
KKYQEWSRKRSIQWEAISEGYKKYKGMDEFKNTFKNIKEPDANEPNANEYLKKHCSKCPCGFN
DMQEITKYTNIGNEAFKQIKEQVDIPAELEDVIYRLKHHEYDKGNDYICNKYKNINVNMKKNND
DTWTDLVKNSSDINKGVLLPPRRKNLFLKIDESDICKYKRDPKLFKDFIYSSAISEVERLKKVYGE
AKTKVVHAMKYSFADIGSIIKGDDMMENNSSDKIGKILGDGVGQNEKRKKWWDMNKYHIWES
MLCGYKHAYGNIAENDRKMLDIPNNDDEHQFLRWFQEWTENFCTKRNELYENMVTACNSAKC
NTSNGSVDKKECTEACKNYANFILIKKKEYQSLNSQYDMNYKETKAEKKESPEYFKDKCNGECS
CLSEYFKDETRWKNPYETLDDTEVKNNCMCKPPPPASNSGRELEFGPEQKLISEEDLNSAVDHH
HHHH

**DBL1x-6ε**

DAAQPARRARRTKLGTDSGTNDPCDRIPPPYGDNDQWKCAIILSKVSEKPENVFVPPRRQRMCIN
NLEKLNVDKIRDKHAFLADVLLTARNEGERIVQNHPDTNSANVCNALERSFADIADIIRGTDLWK

GTNSNLEQNLKQMFAKIRENDKVLQDKYPKDQNYRKLREDWWNANRQKVWEVITCGARSND
LLIKRGWRTSGKSNGDNKLELCRKCGHYEEKVPTKLDYVPQFLRWLTEWIEDFYREKQNLIDD
MERHREECTSEDHKSKEGTSYCSTCKDKCKKYCECVKKWKSEWENQKNKYTELYQQNKNEAS
QKNTSRYDDYVKDFFKKLEANYASLENYIKGDPYFAEYATKLSFILNSADANNPSEKIQKNNDE
VCNCNESGIASVEQEQISDPSSNKACITHSSIKANKKKVCKHVKLGVRENDKDLRVCVIEHTSLS
GVENCCCQDFLRILQENCADNKAGSSSNGACNNKNQEACEKNLEKVLASLTNCYKCDKCKSEQ
SKKNNKNWIWKKSSGKEGGLQKEYANTIGLPPRTQSLCLVVCLDEKGKKTQELKNIRTNSELLK
EWIIAAFHEGKNLKPSHEKKNDDNGKKLCKALEYSFADYGDLIKGTSIWDNEYTKDLELNLQKI
FGKLFRKYIKKNNAAEQDTSYSSLDELRESWWNTNKKYIWLAMKHGAGMNSTTCCGDGSVTG
SGSSCDDIPTIDLIPQYLRFLQEWVEHFCKQRQEKVKPVIENCKSCKESGGTCNGECKTECKNKC
EVYKKFIEDCKGGDGTAGSSWVKRWDQIYKRYSKYIEDAKRNRKAGTKNCGPSSTTNAAENKC
VQSDIDSFFKHLIDIGLTTPSSYLSIVLDDNICGADKAPWTTYTTYTTTEKCNKETDKSKLQQCNT
AVVVNVPSPLGNTPHGYKYACQCKIPTNEETCDDRKEYMNQWSCGSARTMKRGYKNDNYELC
KYNGVDVKPTTVRSNSAKLDDKDVTFFNLFEQWNKEIQYQIEQYMTNTKISCNNEKNVLSRVSD
EAAQPKFSDNERDRNSITHEDKNCKEKCKCYSLWIEKINDQWDKQKDNYNKFQRKQIYDANKG
SQNKKVVSLSNFLFFSCWEEYIQKYFNGDWSKIKNIGSDTFEFLIKKCGNDSGDGETIFSEKLNNA
EKKCKENESTNNKMKSSETSCDCSEPIYIRGCQPKIYDGKIFPGKGGEKQWICKDTIIHGDTNGAC
IPPRTQNLCVGELWDKRYGGRSNIKNDTKESLKQKIKNAIQKETELLYEYHDKGTAIISRNPMKG
QKEKEEKNNDSNGLPKGFCHAVQRSFIDYKNMILGTSVNIYEYIGKLQEDIKKIIEKGTTKQNGK
TVGSGAENVNAWWKGIEGEMWDAVRCAITKINKKQKKNGTFSIDECGIFPPTGNDEDQSVSWF
KEWSEQFCIERLQYEKNIRDACTNNGQGDKIQGDCKRKCEEYKKYISEKKQEWDKQKTKYENK
YVGKSASDLLKENYPECISANFDFIFNDNIEYKTYYPYGDYSSICSCEQVKYYEYNNAEKKNNKL
LCHEKGNDRTWSKKYIKKLENGRTLEGVYVPPRRQQLCLYELFPIIIKNKNDITNAKKELLETLQI
VAEREAYYLWKQYHAHNDATYLAHKKACCAIRGSFYDLEDIIKGNDLVHDEYTKYIDSKLNEIF
DSSNKNDIETKRARTDWWENEAIAVPNIAGANKADPKTIRQLVWDAMQSGVRKAIDEEKEKKK
PNENFPPCMGVQHIGIAKPQFIRWLEEWTNEFCEKYTKYFEDMKSNCNLRKGADDCDDNSNIEC
KKACANYANWLNPKRIEWNGMSNYYNKIYRKSNKESEDGKDYSMIMEPTVIDYLNKRCNGEIN
GNYICCSCKNIGENSASGTVNKKLQKKETQCEDNKGPLDLMNKVLNKMDPKYSEHKMKCTEV
YLEHVEEQLKEIDNAIKDYKLYPLDRCFDDKSKMKVCDLIGDAIGCKHKTKLDELDEWNDVDM
RDPYNKYKGVLIPPRRRQLCFSRIVRGPANLRNLKEFKEEILKGAQSEGKFLGNYYNEDKDKEKA
LEAMKNSFYDYEYIIKGSDMLTNIQFKDIKRKLDRLLEKETNNAEKVDDWWETNKKSIWNAML
CGYKKSGNKIIDPSWCTIPTTETPPQFLRWIKEWGTNVCIQKEEHKEYVKSKCSNVANLGAQESE
SKNCTSEIKKYQEWSRKRSIQWEAISEGYKKYKGMDEFKNTFKNIKEPDANEPNANEYLKKHCS
KCPCGFNDMQEITKYTNIGNEAFKQIKEQVDIPAELEDVIYRLKHHEYDKGNDYICNKYKNINVN
MKKNNDDTWTDLVKNSSDINKGVLLPPRRKNLFLKIDESDICKYKRDPKLFKDFIYSSAISEVERL
KKVYGEAKTKVVHAMKYSFADIGSIIKGDDMMENNSSDKIGKILGDGVGQNEKRKKWWDMNK
YHIWESMLCGYKHAYGNIAENDRKMLDIPNNDDEHQFLRWFQEWTENFCTKRNELYENMVTA
CNSAKCNTSNGSVDKKECTEACKNYANFILIKKKEYQSLNSQYDMNYKETKAEKKESPEYFKD
KCNGECSCLSEYFKDETRWKNPYETLDDTEVKNNCMCKPPPPASNSGRELEFGPEQKLISEEDLN
SAVDHHHHHH

**DBL1X-2IDa (2319)**
DGTDSGTNDPCDRIPPPYGDNDQWKCAIILSKVSEKPENVFVPPRRQRMCINNLEKLNVDKIRDK
HAFLADVLLTARNEGERIVQNHPDTNSANVCNALERSFADIADIIRGTDLWKGTNSNLEQNLKQ
MFAKIRENDKVLQDKYPKDQNYRKLREDWWNANRQKVWEVITCGARSNDLLIKRGWRTSGKS
NGDNKLELCRKCGHYEEKVPTKLDYVPQFLRWLTEWIEDFYREKQNLIDDMERHREECTSEDH
KSKEGTSYCSTCKDKCKKYCECVKKWKSEWENQKNKYTELYQQNKNEASQKNTSRYDDYVK
DFFKKLEANYASLENYIKGDPYFAEYATKLSFILNSADANNPSEKIQKNNDEVCNCNESGIASVE
QEQISDPSSNKACITHSSIKANKKKVCKHVKLGVRENDKDLRVCVIEHTSLSGVENCCCQDFLRIL
QENCADNKAGSSSNGACNNKNQEACEKNLEKVLASLTNCYKCDKCKSEQSKKNNKNWIWKKS
SGKEGGLQKEYANTIGLPPRTQSLCLVVCLDEKGKKTQELKNIRTNSELLKEWIIAAFHEGKNLK

PSHEKKNDDNGKKLCKALEYSFADYGDLIKGTSIWDNEYTKDLELNLQKIFGKLFRKYIKKNNA
AEQDTSYSSLDELRESWWNTNKKYIWLAMKHGAGMNSTTCCGDGSVTGSGSSCDDIPTIDLIPQ
YLRFLQEWVEHFCKQRQEKVKPVIENCKSCKESGGTCNGECKTECKNKCEVYKKFIEDCKGGD
GTAGSSWVKRWDQIYKRYSKYIEDAKRNRKAGTKNCGPSSTTNAAENKCVQSDIDSFFKHLIDI
GLTTPSSYLSIVLDDNICGADKAPWTTYTTYTTTEKCNKETDKSKLQQCNTAVVVNVPSPLGNTP
HGYKYACQCKIPTNEETCDDRKEYMNQWSCGSARTMKRGYKNDNYELCKYNGVDVKPTTVRS
NSAKLDDKDGPENLYFQGNSAVDDYKDHDGDYKDHDIDYKDDDDKLEVLFQGTGIHHHHHH

**ID2b-DBL6ε (2324)**
METDTLLLWVLLLWVPGSTGDGTDDKDVTFFNLFEQWNKEIQYQIEQYMTNTKISCNNEKNVL
SRVSDEAAQPKFSDNERDRNSITHEDKNCKEKCKCYSLWIEKINDQWDKQKDNYNKFQRKQIY
DANKGSQNKKVVSLSNFLFFSCWEEYIQKYFNGDWSKIKNIGSDTFEFLIKKCGNDSGDGETIFSE
KLNNAEKKCKENESTNNKMKSSETSCDCSEPIYIRGCQPKIYDGKIFPGKGGEKQWICKDTIIHGD
TNGACIPPRTQNLCVGELWDKRYGGRSNIKNDTKESLKQKIKNAIQKETELLYEYHDKGTAIISR
NPMKGQKEKEEKNNDSNGLPKGFCHAVQRSFIDYKNMILGTSVNIYEYIGKLQEDIKKIIEKGTT
KQNGKTVGSGAENVNAWWKGIEGEMWDAVRCAITKINKKQKKNGTFSIDECGIFPPTGNDEDQ
SVSWFKEWSEQFCIERLQYEKNIRDACTNNGQGDKIQGDCKRKCEEYKKYISEKKQEWDKQKT
KYENKYVGKSASDLLKENYPECISANFDFIFNDNIEYKTYYPYGDYSSICSCEQVKYYEYNNAEK
KNNKLLCHEKGNDRTWSKKYIKKLENGRTLEGVYVPPRRQQLCLYELFPIIIKNKNDITNAKKEL
LETLQIVAEREAYYLWKQYHAHNDATYLAHKKACCAIRGSFYDLEDIIKGNDLVHDEYTKYIDS
KLNEIFDSSNKNDIETKRARTDWWENEAIAVPNIAGANKADPKTIRQLVWDAMQSGVRKAIDEE
KEKKKPNENFPPCMGVQHIGIAKPQFIRWLEEWTNEFCEKYTKYFEDMKSNCNLRKGADDCDD
NSNIECKKACANYANWLNPKRIEWNGMSNYYNKIYRKSNKESEDGKDYSMIMEPTVIDYLNKR
CNGEINGNYICCSCKNIGENSASGTVNKKLQKKETQCEDNKGPLDLMNKVLNKMDPKYSEHKM
KCTEVYLEHVEEQLKEIDNAIKDYKLYPLDRCFDDKSKMKVCDLIGDAIGCKHKTKLDELDEWN
DVDMRDPYNKYKGVLIPPRRRQLCFSRIVRGPANLRNLKEFKEEILKGAQSEGKFLGNYYNEDK
DKEKALEAMKNSFYDYEYIIKGSDMLTNIQFKDIKRKLDRLLEKETNNAEKVDDWWETNKKSI
WNAMLCGYKKSGNKIIDPSWCTIPTTETPPQFLRWIKEWGTNVCIQKEEHKEYVKSKCSNVANL
GAQESESKNCTSEIKKYQEWSRKRSIQWEAISEGYKKYKGMDEFKNTFKNIKEPDANEPNANEY
LKKHCSKCPCGFNDMQEITKYTNIGNEAFKQIKEQVDIPAELEDVIYRLKHHEYDKGNDYICNKY
KNINVNMKKNNDDTWTDLVKNSSDINKGVLLPPRRKNLFLKIDESDICKYKRDPKLFKDFIYSSA
ISEVERLKKVYGEAKTKVVHAMKYSFADIGSIIKGDDMMENNSSDKIGKILGDGVGQNEKRKK
WWDMNKYHIWESMLCGYKHAYGNIAENDRKMLDIPNNDDEHQFLRWFQEWTENFCTKRNEL
YENMVTACNSAKCNTSNGSVDKKECTEACKNYANFILIKKKEYQSLNSQYDMNYKETKAEKKE
SPEYFKDKCNGECSCLSEYFKDETRWKNPYETLDDTEVKNNCMCKPPPPASNSGRELEFGPENL
YFQGNSAVDDYKDHDGDYKDHDIDYKDDDDKLEVLFQGTGIHHHHHH

**DBL3X-DBL6ε (2185)**
DGTMKSSETSCDCSEPIYIRGCQPKIYDGKIFPGKGGEKQWICKDTIIHGDTNGACIPPRTQNLCV
GELWDKRYGGRSNIKNDTKESLKQKIKNAIQKETELLYEYHDKGTAIISRNPMKGQKEKEEKNN
DSNGLPKGFCHAVQRSFIDYKNMILGTSVNIYEYIGKLQEDIKKIIEKGTTKQNGKTVGSGAENV
NAWWKGIEGEMWDAVRCAITKINKKQKKNGTFSIDECGIFPPTGNDEDQSVSWFKEWSEQFCIE
RLQYEKNIRDACTNNGQGDKIQGDCKRKCEEYKKYISEKKQEWDKQKTKYENKYVGKSASDLL
KENYPECISANFDFIFNDNIEYKTYYPYGDYSSICSCEQVKYYEYNNAEKKNNKLLCHEKGNDRT
WSKKYIKKLENGRTLEGVYVPPRRQQLCLYELFPIIIKNKNDITNAKKELLETLQIVAEREAYYL
WKQYHAHNDATYLAHKKACCAIRGSFYDLEDIIKGNDLVHDEYTKYIDSKLNEIFDSSNKNDIET
KRARTDWWENEAIAVPNIAGANKADPKTIRQLVWDAMQSGVRKAIDEEKEKKKPNENFPPCMG
VQHIGIAKPQFIRWLEEWTNEFCEKYTKYFEDMKSNCNLRKGADDCDDNSNIECKKACANYAN
WLNPKRIEWNGMSNYYNKIYRKSNKESEDGKDYSMIMEPTVIDYLNKRCNGEINGNYICCSCKN
IGENSASGTVNKKLQKKETQCEDNKGPLDLMNKVLNKMDPKYSEHKMKCTEVYLEHVEEQLK
EIDNAIKDYKLYPLDRCFDDKSKMKVCDLIGDAIGCKHKTKLDELDEWNDVDMRDPYNKYKG

VLIPPRRRQLCFSRIVRGPANLRNLKEFKEEILKGAQSEGKFLGNYYNEDKDKEKALEAMKNSFY
DYEYIIKGSDMLTNIQFKDIKRKLDRLLEKETNNAEKVDDWWETNKKSIWNAMLCGYKKSGNK
IIDPSWCTIPTTETPPQFLRWIKEWGTNVCIQKEEHKEYVKSKCSNVANLGAQESESKNCTSEIKK
YQEWSRKRSIQWEAISEGYKKYKGMDEFKNTFKNIKEPDANEPNANEYLKKHCSKCPCGFNDM
QEITKYTNIGNEAFKQIKEQVDIPAELEDVIYRLKHHEYDKGNDYICNKYKNINVNMKKNNDDT
WTDLVKNSSDINKGVLLPPRRKNLFLKIDESDICKYKRDPKLFKDFIYSSAISEVERLKKVYGEAK
TKVVHAMKYSFADIGSIIKGDDMMENNSSDKIGKILGDGVGQNEKRKKWWDMNKYHIWESML
CGYKHAYGNIAENDRKMLDIPNNDDEHQFLRWFQEWTENFCTKRNELYENMVTACNSAKCNT
SNGSVDKKECTEACKNYANFILIKKKEYQSLNSQYDMNYKETKAEKKESPEYFKDKCNGECSCL
SEYFKDETRWKNPYETLDDTEVKNNGPENLYFQGNSAVDDYKDHDGDYKDHDIDYKDDDDKL
EVLFQGISVPSIPPDVSGFSIGIHHHHHH

**DBL4ε-DBL6ε (2149)**
DQVKYYEYNNAEKKNNKLLCHEKGNDRTWSKKYIKKLENGRTLEGVYVPPRRQQLCLYELFPII
IKNKNDITNAKKELLETLQIVAEREAYYLWKQYHAHNDATYLAHKKACCAIRGSFYDLEDIIKG
NDLVHDEYTKYIDSKLNEIFDSSNKNDIETKRARTDWWENEAIAVPNIAGANKADPKTIRQLVW
DAMQSGVRKAIDEEKEKKKPNENFPPCMGVQHIGIAKPQFIRWLEEWTNEFCEKYTKYFEDMKS
NCNLRKGADDCDDNSNIECKKACANYANWLNPKRIEWNGMSNYYNKIYRKSNKESEDGKDYS
MIMEPTVIDYLNKRCNGEINGNYICCSCKNIGENSASGTVNKKLQKKETQCEDNKGPLDLMNKV
LNKMDPKYSEHKMKCTEVYLEHVEEQLKEIDNAIKDYKLYPLDRCFDDKSKMKVCDLIGDAIG
CKHKTKLDELDEWNDVDMRDPYNKYKGVLIPPRRRQLCFSRIVRGPANLRNLKEFKEEILKGAQ
SEGKFLGNYYNEDKDKEKALEAMKNSFYDYEYIIKGSDMLTNIQFKDIKRKLDRLLEKETNNAE
KVDDWWETNKKSIWNAMLCGYKKSGNKIIDPSWCTIPTTETPPQFLRWIKEWGTNVCIQKEEHK
EYVKSKCSNVANLGAQESESKNCTSEIKKYQEWSRKRSIQWEAISEGYKKYKGMDEFKNTFKNI
KEPDANEPNANEYLKKHCSKCPCGFNDMQEITKYTNIGNEAFKQIKEQVDIPAELEDVIYRLKHH
EYDKGNDYICNKYKNINVNMKKNNDDTWTDLVKNSSDINKGVLLPPRRKNLFLKIDESDICKYK
RDPKLFKDFIYSSAISEVERLKKVYGEAKTKVVHAMKYSFADIGSIIKGDDMMENNSSDKIGKIL
GDGVGQNEKRKKWWDMNKYHIWESMLCGYKHAYGNIAENDRKMLDIPNNDDEHQFLRWFQE
WTENFCTKRNELYENMVTACNSAKCNTSNGSVDKKECTEACKNYANFILIKKKEYQSLNSQYD
MNYKETKAEKKESPEYFKDKCNGECSCLSEYFKDETRWKNPYETLDDTEVKNNGPEQKLISEED
LNSAVDHHHHHH

**Table S1**. Amino acid sequences of constructs described in the text as defined in Figure 1A.

| | | |
|---|---|---|
| DBL1x-2IDa (flag) | GGGCCCGAGAACCTGTAC | GTCCTTGTCGTCCAGCTTAG |
| DBL1x-2IDb (flag) | GGGCCCGAGAACCTGTAC | CTTGTTGTTGGTGGACTCGTTTT |
| ID2b-DBL6ε (flag) | GACGACAAGGACGTCACC | GGTACCGTCACCAGTGGA |
| DBL3x-DBL6ε flag | ATGAAGTCCTCTGAAACCTCCT | GTCACCAGTGGAACCTGG |
| DBL4ε-DBL6ε myc | CAAGTGAAGTACTACGAGTAC | GTCACCAGTGGAACCTGG |
| Removal of N-terminal cloning artefact from vector | GTC ACC AGT GGA ACC TGG AAC CGGT | GGT ACC GAA TTC GGG CCC |
| Truncation of C-Terminus | GAACAAAAACTCATCTCAGAAG | GTTGTTCTTGACTTCGGTG |
| Removal of ApaI site and cmyc tag-introduction of EcoRV site | TCCATCATCATCATCATCATTGAG TTTAAACC | TATCGGTACCGTCACCAGTG |

| | |
|---|---|
| GBLOCK Amino acids 1-58 + KpnI | AAGTCCTCTATCGCCAACAAGATCGAAGCCTACCTGGGAGC CAAGTCTGACGACTCTAAGATCGACCAGTCTCTGAAGGCCG ACCCCTCTGAAGTCCAGTACTACGGATCTGGTGGCGACGGT TACTACCTGAGGAAGAACATCTGCAAGATCACCGTCAACCA CGGTACCGACTCTGGG |
| GBLOCK EcoRI+ApaI site + TEV protease + 3XFLAG | GAATTCGGGCCCGAGAACCTGTACTTCCAAGGCAATAGCGC CGTCGACGACTACAAAGACCATGACGGTGATTATAAAGATC ATGACATCGATTACAAGGATGACGATGACAAGCTGGAAGTT CTGTTCCAGGGGACCGGG |

**Table S2** Primers used to generate constructs.

| | NTS-DBL6ε | DBL3x-DBL6ε | DBL4ε-DBL6ε | DBL1x-ID2a |
|---|---|---|---|---|
| Experimental Date | 11-01-2017 | 11-01-2017 | 07-08-2018 | 03-05-2018 |
| Location | APS | APS | NSLSII | NSLSII |
| Beamline/instrument | 18-ID | 18-ID | 16-ID | 16-ID |
| instrument | Pilatus 1M | Pilatus 1M | Pilatus 1M (SAXS) and 3K (WAXS) | Pilatus 1M (SAXS) and 3K (WAXS) |
| SEC flow rate (mls/min) | 0.5 | 0.5 | 0.5 | 0.5 |
| Cell temperature | RT | RT | RT | RT |
| Protein concentration (mg/ml) | | | 10.2 | 7.5 |
| Number of residues, including tags | 2671 | 1483 | 1095 | 1016 |
| Molecular weight (calculated) | 309845 | 173423 | 128802 | 116489 |

**Table S3** Experimental details for SAXS data collection

| | Residue number | Template | Matched residues | % Identity (similarity) | Residues modeled | zDOPE* |
|---|---|---|---|---|---|---|
| DBL1x | 10-388 | 2YK0 | 31-434 | 30 (54) | 10-410 | -0.42 (-0.30) |
| ID1 | 372-580 | NT | | | | |
| DBL2x | 581-906 | 2XU0_A | 132-447 | 23 (49) | 569-905 | 0.04 (0.08) |
| ID2a | 907-1078 | NT | | | | |
| ID2b | 1079-1149 | 4P1T | 1833-1905 | 22 (52) | | |
| | 1150-1216 | NT | | | | |
| DBL3x-DBL4ε | 1207-1944 | 4P1T | 1215-1949 | 83 (89) | 1205-1944 | -0.55 (-0.55) |
| | 1965-1978 | NT | | | | |
| DBL5ε | 1979-2290 | 3VUV_A | 41-446 | 31 (54) | 1979-2283 | -0.55 (-0.51) |
| | 2291-2332 | NT | | | | |
| DBL6ε | 2333-2364 | 2WAU_A | 2333-2634 | 99 (99) | 2350-2634 | -1.89 (-1.88) |

**Table S4**. Regions modelled using Modeller and templates used.

NT = no template; GA341 = 1 in all cases
